# Zebrafish knockout models of *atxn1a*, *atxn1b*, and *atxn1l* reveal distinct and shared phenotypic and transcriptomic alterations

**DOI:** 10.64898/2026.03.05.709450

**Authors:** Anwarul Karim, Pramuk Keerthisinghe, Sreeja Sarasamma, Nicholas A. Ciaburri, Mabel Guerra Giraldez, Kylan Naidoo, James P. Orengo

## Abstract

Spinocerebellar ataxia type 1 (SCA1) is a progressive neurodegenerative disorder caused by polyglutamine expansion in ATXN1, yet the normal physiological roles of ATXN1 and its paralog ATXN1L remain incompletely understood. To define these roles, we generated the first zebrafish knockouts (KOs) of the three ataxin-1 family genes, *atxn1a*, *atxn1b*, and *atxn1l*, using CRISPR/Cas9. These mutants reveal distinct and shared developmental, behavioral, and transcriptomic alterations. All KOs showed reduced early survival and mild larval growth deficits, indicating essential developmental functions. Behavioral assays revealed distinct paralog-specific effects: *atxn1a* KO larvae exhibited a unique light-dependent locomotor deficit, whereas *atxn1b* and *atxn1l* KOs displayed global hypoactivity. Adult behavioral assessment revealed a gradient of phenotypic severity: *atxn1a* KOs displayed the earliest and most pronounced alterations in vertical tank exploration and the greatest impairment in swim-tunnel performance, followed by *atxn1b* and then *atxn1l* mutants. To define molecular mechanisms underlying these phenotypes, we performed RNA-seq at 5 days post-fertilization and identified unique and shared differentially expressed genes across the three KO lines. Shared transcriptomic signatures highlighted suppression of leukotriene-biosynthetic pathways and diminished innate-immune pathways; suggesting that ATXN1-family genes influence neuroimmune signaling during early development. Weighted gene co-expression network analysis identified distinct KO-associated gene modules, including a phototransduction-enriched module strongly correlated with *atxn1a* KO status, offering a mechanistic link to its light-dependent locomotor phenotype. Together, these findings establish a comprehensive assessment of zebrafish models that reveal both shared core functions and specialized roles of ATXN1-family genes in development, neuroimmune regulation, sensorimotor behavior, and retinal signaling.

## Introduction

Spinocerebellar ataxia type 1 (SCA1) is a progressive, autosomal dominant neurodegenerative disorder caused by an expanded CAG repeat (polyglutamine, polyQ) in the coding region of the *ATXN1* (ataxin-1) gene (Zoghbi & Orr, 1995). It affects approximately 1-2 in 100,000 individuals and is characterized by progressive motor incoordination or ataxia (Tejwani & Lim, 2020). Symptom onset typically occurs in the second to third decade of life, with disease progression spanning 10 to 30 years and there is no curative treatment available. In addition to ataxia, patients may present with dysarthria, dysphagia, respiratory dysfunction, spasticity, muscle weakness and wasting, ophthalmoplegia, cognitive impairment, and sensory neuropathy (Olmos et al., 2022). The hallmark pathological finding is the degeneration of cerebellar Purkinje cells. However, degeneration in other regions, including the brainstem and spinal cord, is also present (Tejwani & Lim, 2020). Previous studies from mouse model of SCA1 suggested that the neurotoxicity associated with mutant ATXN1 protein results from its incorporation into endogenous complexes containing Capicua (Lam et al., 2006). Overexpression of ATXN1L (ataxin-1-like), a paralog of ATXN1, suppresses neuropathology by displacing mutant ATXN1 from its complex with Capicua (Carrell et al., 2022; Crespo-Barreto et al., 2010; Mizutani et al., 2005).

Beyond SCA1, variants in *ATXN1* have been reported to be associated with an increased risk of Alzheimer’s disease (Suh et al., 2019). Studies in mice showed that loss of *Atxn1* elevates BACE1 protein expression and Aβ pathology, suggesting a potential contribution to Alzheimer’s disease pathogenesis (Suh et al., 2019). Furthermore, *ATXN1* has also been identified as a susceptibility locus for multiple sclerosis (International Multiple Sclerosis Genetics Consortium et al., 2019). ATXN1 exerts a protective role against autoimmune demyelination in a preclinical mouse model of multiple sclerosis, and its loss of function results in increased demyelination, axonal degeneration, and oligodendrocyte loss (Didonna et al., 2020; Talukdar et al., 2025). Furthermore, mice lacking *Atxn1* exhibit learning deficits as well as transcriptomic and proteomic alterations (Crespo-Barreto et al., 2010; Goold et al., 2007; Matilla et al., 1998; Sánchez et al., 2016). On the other hand, loss of *Atxn1l* in mice results in reduced body size, developmental abnormalities, and increased mortality, phenotypes that are further exacerbated in *Atxn1*/*Atxn1l* double knockouts, indicating partial functional redundancy between these genes (Lee et al., 2011). Together, these findings demonstrate that ATXN1 and ATXN1L have diverse biological roles extending well beyond SCA1. However, despite substantial progress in understanding SCA1 pathology, the normal physiological functions of ATXN1 and ATXN1L remain incompletely understood. Elucidating these functions and the consequences of their perturbation is therefore crucial for the development of successful therapeutic strategies, especially because some current approaches interfere with the expression of the wild-type (WT) protein (Kerkhof et al., 2023).

Zebrafish have emerged as a valuable model for studying gene functions, disease pathogenesis, and high-throughput drug screening in neurodegenerative diseases (Chia et al., 2022; Saleem & Kannan, 2018; Sarasamma et al., 2023; Sherman et al., 2022; Srivastava et al., 2025). The zebrafish genome contains two *ATXN1* orthologs, *atxn1a* and *atxn1b*, and an *ATXN1L* ortholog, *atxn1l* (Carlson et al., 2009; Vauti et al., 2021). Protein sequence analysis revealed that zebrafish Atxn1a and Atxn1b share 46% and 37% identity, respectively, with human ATXN1, and zebrafish Atxn1l shares 40% identity with human ATXN1L (Carlson et al., 2009). In particular, the N-terminal region, AXH domain, and C-terminal region of human ATXN1 and ATXN1L exhibit significant conservation across the three zebrafish homologs. Furthermore, the nuclear localization sequence in human ATXN1 protein, centered around K772 and S776, is also conserved in zebrafish Atxn1a and Atxn1b. An important distinction is that the CAG repeat found in human *ATXN1* gene does not exist in zebrafish *atxn1a*, *atxn1b*, and *atxn1l*; this absence is also observed in mouse *Atxn1* and *Atxn1l* (Carlson et al., 2009; Vauti et al., 2021). A whole-mount *in situ* hybridization study on zebrafish embryos and larvae revealed strong central nervous system expression of these three genes, including the cerebellum (Vauti et al., 2021). This suggests that the *atxn1*-family may play an important role in zebrafish neurodevelopment.

In a recent study, a stable transgenic zebrafish model of SCA1 was established (Elsaey et al., 2021). The authors expressed cDNA encoding human ATXN1 that either contains a non-pathological polyQ stretch of 30 glutamine residues (Atx1[30Q]) interrupted by two histidine residues or a patient-derived pathological polyQ stretch of 82 non-interrupted glutamine residues (Atx1[82Q]). By using the regulatory element cpce, a small genomic sequence upstream of the zebrafish gene carbonic anhydrase 8 (*ca8*), they were able to express the transgene specifically in Purkinje cells. Atx1[82Q] fish exhibited progressive degeneration of Purkinje cells, beginning in the larval stage and becoming more prominent in young adults. Notably, the degeneration was more pronounced in the rostral region compared to the caudal regions of the Purkinje cell population, highlighting differential vulnerability even within the Purkinje cell population. While this SCA1 model is expected to significantly contribute to understanding the mechanism of SCA1 caused by pathological polyQ ATXN1, there is still a lack of models to study the normal functions of *ATXN1* and *ATXN1L* orthologs in zebrafish. Developing a range of complementary models suited to investigate different aspects, such as the normal functions of the relevant genes and the mechanisms of disease resulting from the disease-specific mutation, is essential for a comprehensive understanding of the disease and the development of potential therapies. To address this, we developed zebrafish knockout (KO) models of *atxn1a*, *atxn1b*, and *atxn1l* using CRISPR-Cas9. We used these models to investigate the phenotypic alterations in both larval and adult stages and the transcriptional alterations at the larval stage. We found that the KO of these genes resulted in varying effects on both phenotype and transcriptome. These models are a robust resource for dissecting ATXN1 biology, understanding the consequences of gene-lowering therapeutic strategies, and accelerating cross-species mechanistic discovery in SCA1 and ATXN1-related neurodegeneration.

## Results

### Generation and validation of knock-out mutations in *atxn1a*, *atxn1b*, and *atxn1l*

To investigate the consequences of *atxn1a*, *atxn1b*, and *atxn1l* KO in zebrafish, we used CRISPR/Cas9 technology to introduce frameshift mutations in each gene separately (Fig. 1). WT AB zebrafish embryos were injected at the 1-cell stage into the cytoplasm with Cas9 protein and an sgRNA (Fig. 1C) targeting the respective genes. The first coding exon of each gene was selected as the target (Fig. 1B). Injected embryos (F_0_) were grown to adulthood and outcrossed to WT fish to generate the F_1_ generation. Fin clipping of F_1_ fish was performed to identify heterozygous fish with frameshift mutations predicted to be disruptive of protein expression. Selected F_1_ fish (14 bp deletion in *atxn1a*, 1 bp deletion and 6 bp substitution in *atxn1b*, and a 4 bp deletion in *atxn1l*) were crossed to generate F_2_ homozygous fish, which were raised to maintain the homozygous lines and to conduct experiments. The mutations were validated by Sanger sequencing as well as RNA-seq (Fig. 1B).

**Fig. 1.**
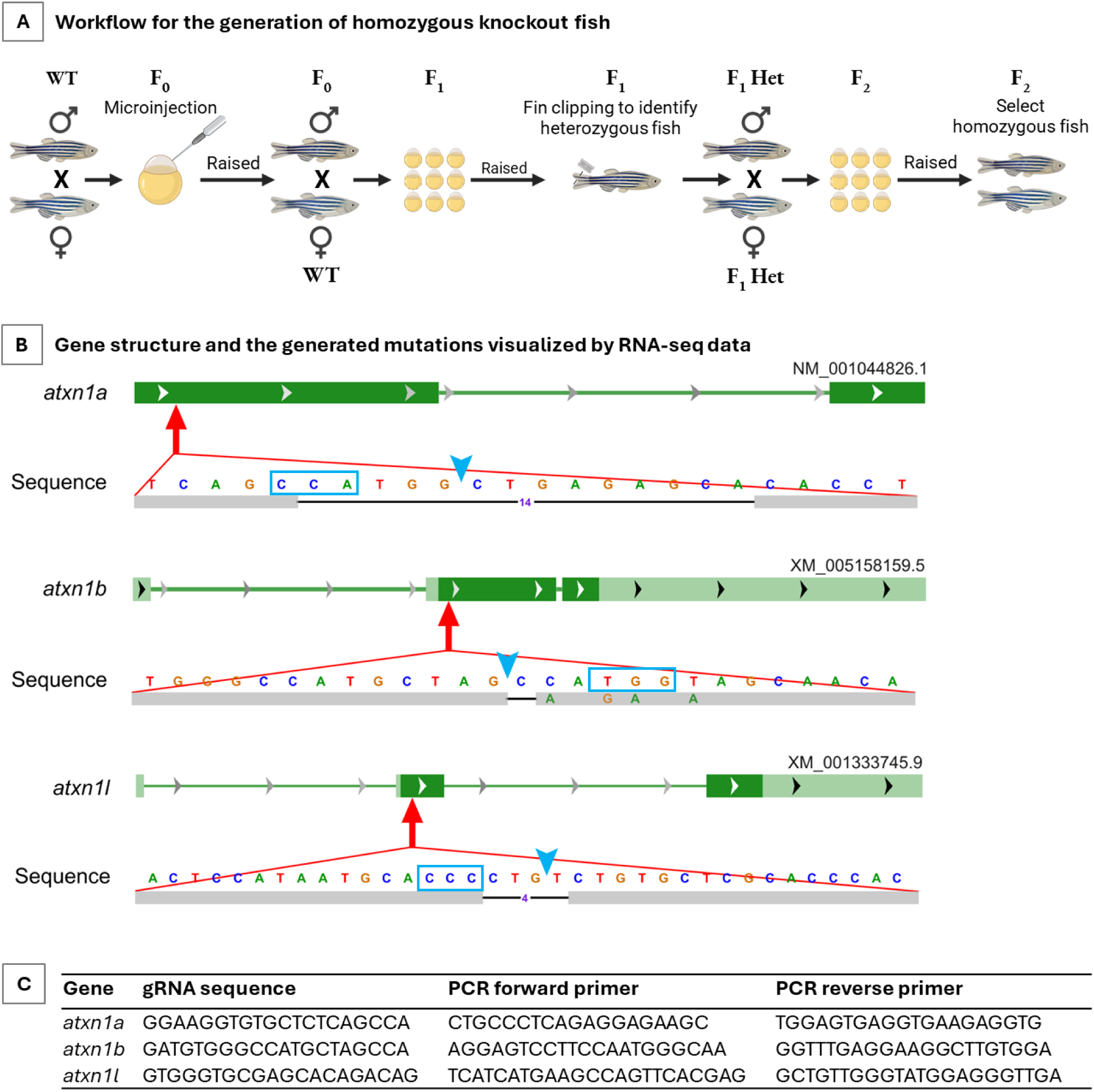
Generation and validation of frameshift mutations in *atxn1a*, *atxn1b*, and *atxn1l* in zebrafish. **A.** Workflow for the generation of homozygous knockout fish using CRISPR/Cas9 technology. **B.** Gene structure of *atxn1a*, *atxn1b*, and *atxn1l*. The first coding exon of each gene was targeted using CRISPR/Cas9. The resulting mutations in each gene and the flanking sequences are shown from RNA-seq reads visualized in the Integrative Genomics Viewer. The PAM sequence for each gene is shown in a blue box, and the cleavage site is indicated by a blue arrowhead. **C.** Table listing the gRNA sequences used to target the genes and the PCR primers flanking the target sites that were used for genotyping.

### Survival, hatching rate, and body length of the KO larvae

After generating homozygous KO lines of *atxn1a*, *atxn1b*, and *atxn1l*, we investigated larval survival, hatching rate, and body length. For survival analysis, in each experiment we used two Petri dishes, each containing 50 viable embryos, selected by unaided visual inspection, at 2 hours post-fertilization. Live embryos were counted on days 1, 2, 3, and 5 post-fertilization. The experiment was repeated 10 times; therefore, the total starting number of embryos for each fish line assessed was 1000. Survival analysis using the log-rank (Mantel–Cox) test, comparing WT with each homozygous KO group separately, showed decreased survival in all KO lines compared to WT, with most deaths occurring within the first day post-fertilization (dpf) (Fig. 2A). Separate analysis of the same data showing the percentage of surviving embryos at 1 dpf revealed significantly lower survival rates in all KO groups compared to WT (mean survival rates: 84.8 ± 11.3%, 59.6 ± 12.5%, 59.2 ± 17.1%, and 66.1 ± 15.9% in WT, *atxn1a* KO, *atxn1b* KO, and *atxn1l* KO, respectively) (Fig. 2B). From the same Petri dishes, we also counted the total number of hatched embryos at 2 and 3 dpf. As shown in Fig. 2C, *atxn1a* KO embryos exhibited significantly lower hatching rate (1.5 ± 3.9%) at 2 dpf compared to WT (24 ± 23.8%), whereas nearly all embryos in all fish lines had hatched by 3 dpf. Next, we assessed body size of larva for each KO. At 5 dpf, images of 10 larvae per group per experiment were taken using a microscope, and this was repeated across four independent experiments (n = 40 per group). Larval length from head to tail was systematically measured as shown in Fig. 2E. The analysis revealed that all KO groups had slightly reduced but statistically significant mean body length compared to WT, with the *atxn1l* KO group showing the shortest mean length (Fig. 2F). This observation is consistent with previous reports of reduced body size in *Atxn1l*^−/−^ mice (Lee et al., 2011).

**Fig. 2.**
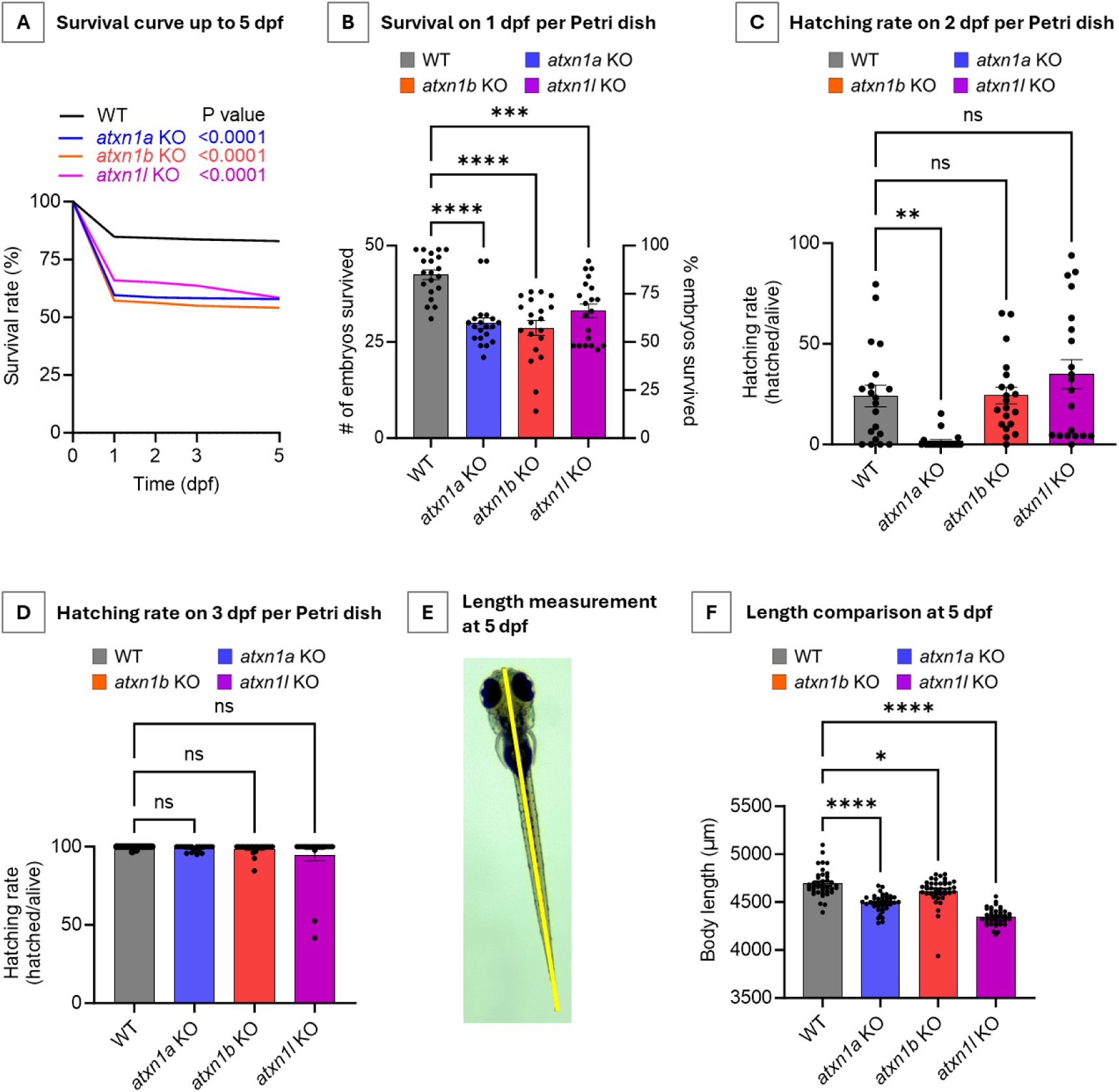
Characterization of homozygous KO embryos and larvae up to 5 dpf. **A.** Survival curves of WT, *atxn1a*, *atxn1b*, and *atxn1l* KOs up to 5 dpf. Survival of each KO was compared separately with WT using the log-rank (Mantel–Cox) test, and the P values are shown in the figure. Data were acquired from 10 independent experiments. Each experiment started with two Petri dishes containing 50 embryos per dish (100 embryos per experiment) for each group. n = 1000 embryos per group. **B.** Survival at 1 dpf. Data are from the same experiments shown in panel **A**. Each dot represents the number (or proportion) of embryos that survived in each Petri dish by 1 dpf. All KO lines show reduced survival compared with WT. *** P < 0.001, **** P < 0.0001; ordinary one-way ANOVA with Šídák’s multiple comparisons test. **C, D.** Hatching rate at 2 dpf (**C**) and 3 dpf (**D**). Each dot represents the number of hatched embryos divided by the total number of live embryos at the time of observation. Data were collected from the same Petri dishes used in panel **A** (10 experiments, 20 Petri dishes per group). At 2 dpf, *atxn1a* KO embryos show reduced hatching rate compared with WT. ** P < 0.01; ns, not significant; ordinary one-way ANOVA with Šídák’s multiple comparisons test. **E, F.** Larval length (head to tail) at 5 dpf was measured (**E**). All KO larvae show reduced body length compared with WT (**F**). * P < 0.05, **** P < 0.0001; ordinary one-way ANOVA with Šídák’s multiple comparisons test.

### *atxn1a* KO larvae exhibit light-dependent locomotor phenotype at 5 dpf

Locomotor activity tracking of zebrafish larva under light and dark conditions has commonly been used as an easily accessible phenotypic readout in various studies (Tuz-Sasik et al., 2022). To characterize the locomotor activity of our homozygous KO larvae, we used the DanioVision system for high-throughput larval tracking at 5 dpf. We designed two light–dark paradigms as follows: (i) 40 minutes of dark habituation, 10 minutes of light, and 10 minutes of dark, with this light–dark cycle repeated four times (hereafter referred to as the “short light–dark paradigm”); and (ii) 1 hour of dark habituation, 1 hour of light, and 1 hour of dark, with this light–dark cycle repeated twice (hereafter referred to as the “long light–dark paradigm”) (Fig. 3A). Each experiment was repeated five times for each KO group using different batches of larvae. In each experiment, a 96-well plate was used, containing 48 WT larvae and 48 KO larvae. All experiments were initiated at the same time of day, between 12:00 pm and 1:00 pm, and required a total of 7 hours to complete, during which the short light–dark paradigm was performed first, followed by the long light–dark paradigm.

**Fig. 3.**
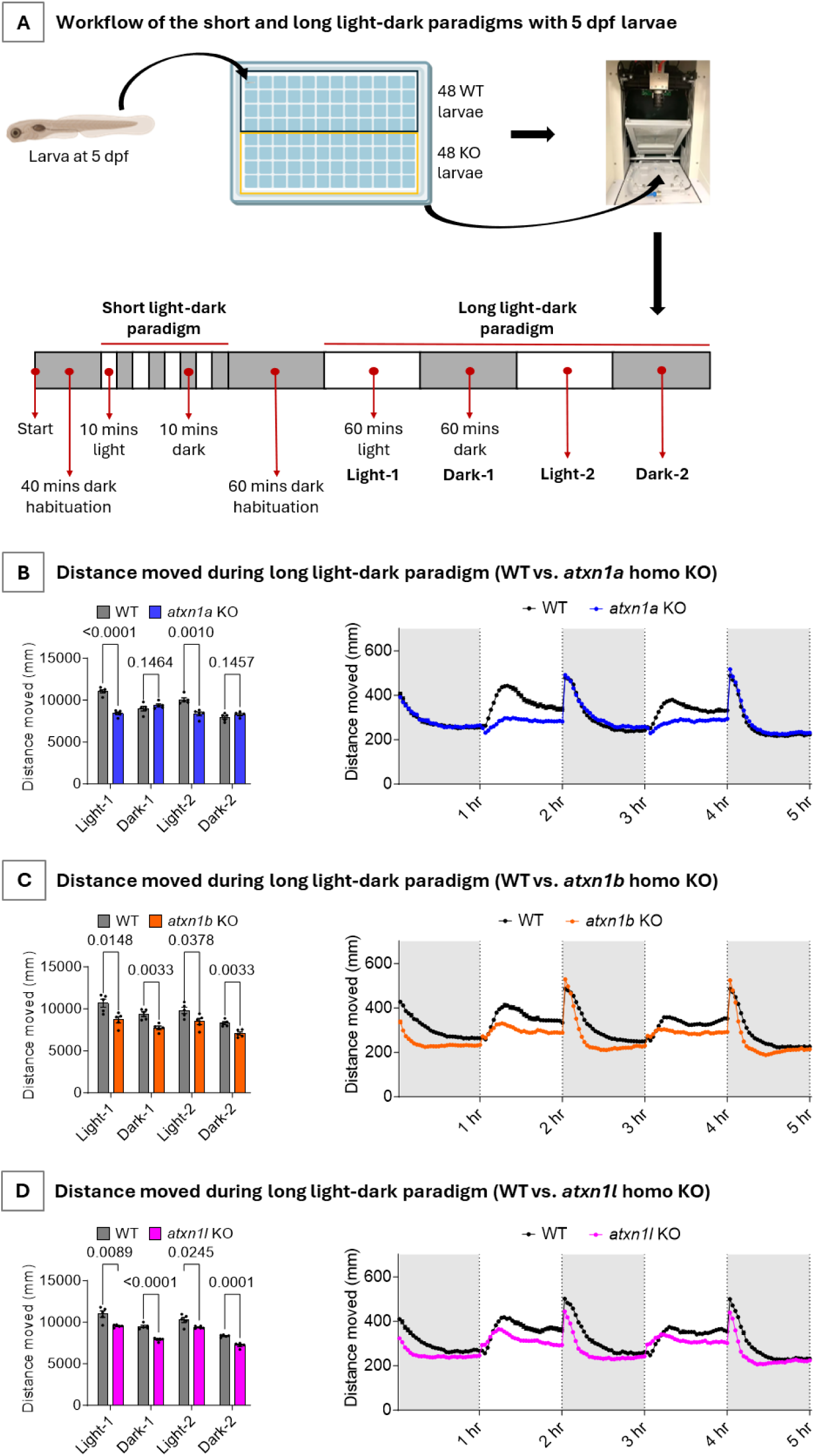
Assessment of locomotor activity of homozygous KO larvae at 5 dpf. **A.** Workflow of locomotor activity recording using DanioVision to analyze the response of 5 dpf larvae to light–dark paradigms. The 7-hour experiment consisted of an initial habituation period, followed by the short light–dark paradigm, a second habituation period, and then the long light–dark paradigm. **B–D.** Distance moved during the long light–dark paradigm. Data represent combined results from five independent experiments for each KO line. In each experiment, 48 WT larvae and 48 KO larvae were used. Thus, for each KO line, a total of 240 WT larvae and 240 KO larvae were used. The left panels show the cumulative distance moved during each 1-hour light or dark phase of the long light–dark paradigm. Each dot represents the mean distance moved in one experiment. Statistical analysis was performed using multiple unpaired t tests with FDR correction by the two-stage step-up method of Benjamini, Krieger, and Yekutieli. The q values are shown in the graph. The right panels show the distance moved in each 2-min interval throughout the long light–dark paradigm. The results show that *atxn1a* KO larvae (**B**) exhibit reduced locomotor activity specifically during the light phase, whereas *atxn1b* (**C**) and *atxn1l* (**D**) KO larvae show a generalized reduction in locomotor activity during both light and dark phases.

The recorded videos were analyzed with EthoVision XT 13 to calculate the distance traveled by each larva. The short paradigm showed an expected striking drop in movement immediately after the transition from the dark to light phase, and an increase in movement immediately after the transition from the light to dark phase (Supplementary Fig. 1-3). This phenomenon was observed in WT as well as all KO groups. However, the short paradigm failed to reveal reproducible patterns in the KO groups that were clearly distinguishable from the WT group across multiple experiments. In contrast, the long light–dark paradigm revealed consistent and robust phenotypic differences between KO and WT larvae (Supplementary Fig. 1-3). Both *atxn1b* and *atxn1l* KO larvae traveled shorter distances than WT larvae under both light and dark conditions (Fig 3C and 3D). Surprisingly, *atxn1a* KO larvae exhibited reduced movement only during the light phase, while traveling distances comparable to WT larvae during the dark phase (Fig. 3B). This suggests a unique, light-dependent locomotor phenotype specific to *atxn1a* KO larvae, whereas the other two KO lines exhibit a more generalized reduction in locomotor activity.

### KO of *atxn1* family members exhibit behavioral deficits in adulthood, as assessed by novel tank and swim tunnel assays

We assessed adult swimming behavior using two commonly employed assays: novel static tank and swim tunnel. For analyzing swimming behavior in a novel tank, we followed our recently described protocol (Keerthisinghe et al., 2026). Briefly, we filled clear aquatic habitat tanks (20 cm L × 12 cm H × 9 cm W; ≈ 1.8 L) with fresh system water (Fig. 4A), placed a single adult zebrafish in the tank, acclimated the fish for 5 minutes, and then started recording video of the fish exploring the tank. Videos of 10 minutes duration were analyzed with DeepLabCut (Mathis et al., 2018) using a pretrained model (Keerthisinghe et al., 2026) (Fig. 4B). Zone occupancy (top, middle, and bottom zones) by each fish over a 10-minute recording was calculated. The experiment was performed for each KO line together with WT controls at three age time points; 8, 12, and 16 months. At these time-points, we found that WT fish spent most of the time in the bottom zone of the tank and the least time in the top zone (Fig. 4C, 4G, 4I, and 4J). However, *atxn1a* homozygous KO fish spent significantly more time in the top zone and less time in the bottom zone compared to WT at all time points (Fig. 4D, 4G, 4I, and 4J). For *atxn1b* homozygous KO fish, a trend toward spending less time in the bottom zone and more time in the middle zone compared to WT was observed; this difference was not statistically significant at 8 months (Fig. 4E and 4G) but became statistically significant at 12 and 16 months (Fig. 4I and 4J), indicating a progressive phenotype with a delayed onset in *atxn1b* KO fish compared to *atxn1a* KO fish. Lastly, *atxn1l* homozygous KO fish did not show any statistically significant differences in zone occupancy compared to WT at 8 or 12 months (Fig. 4F, 4G, and 4I), but at 16 months they spent less time in the bottom zone and more time in the middle zone compared to WT, indicating that this phenotype develops even more slowly in *atxn1l* KO fish. We also conducted this experiment with 8-month-old heterozygous KO fish for all three paralogs and found no difference compared to WT in terms of zone occupancy (Fig. 4H).

**Fig. 4.**
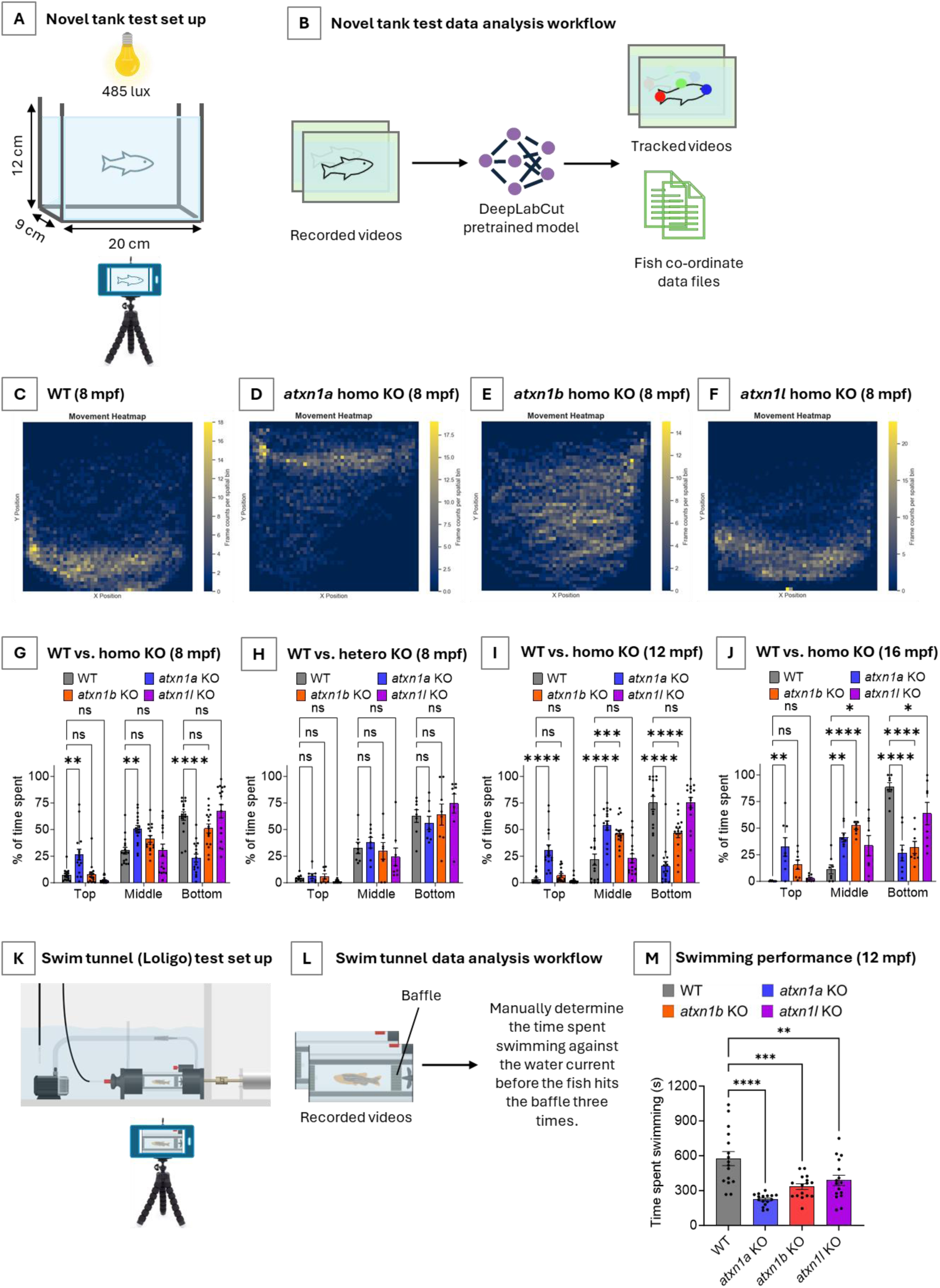
Phenotypic assessment of adult fish. **A.** Novel tank test setup for recording the activity of adult fish when placed in a novel tank. **B.** Workflow for tracking fish in recorded videos using DeepLabCut. **C–F.** Representative heatmaps of 8-month-old WT (**C**), *atxn1a* homozygous KO (**D**), *atxn1b* homozygous KO (**E**), and *atxn1l* homozygous KO (**F**) fish showing zone occupancy during a 10-min recording. **G.** Time spent in the top, middle, and bottom zones of novel tank by WT and homozygous KO fish at 8 mpf. ** P < 0.01; **** P < 0.0001; ns, not significant; ordinary two-way ANOVA with Dunnett’s multiple comparisons test. n = 16 (8 male + 8 female) per group. **H.** Time spent in the top, middle, and bottom zones of novel tank by WT and heterozygous KO fish at 8 mpf. ns, not significant; ordinary two-way ANOVA with Dunnett’s multiple comparisons test. n = 8 (4 male + 4 female) per group. **I.** Time spent in the top, middle, and bottom zones of novel tank by WT and homozygous KO fish at 12 mpf. *** P < 0.001; **** P < 0.0001; ns, not significant; ordinary two-way ANOVA with Dunnett’s multiple comparisons test. n = 16 (8 male + 8 female) per group. **J.** Time spent in the top, middle, and bottom zones of novel tank by WT and homozygous KO fish at 16 mpf. * P < 0.05; ** P < 0.01; **** P < 0.0001; ns, not significant; ordinary two-way ANOVA with Dunnett’s multiple comparisons test. n = 8 (4 male + 4 female) per group. **K.** Swim tunnel test setup for recording swimming performance of adult fish against water current. **L.** Data analysis workflow for videos recorded during the swim tunnel test. **M.** Swimming performance of 12 mpf fish, measured as the time spent swimming against increasing speed of water current before the fish contacted the baffle three times. ** P < 0.01; *** P < 0.001; **** P < 0.0001; ordinary one-way ANOVA with Šídák’s multiple comparisons test. n = 16 (8 male + 8 female) per group.

Given that the behavioral alterations observed in the novel tank test showed gradual development across different KO lines between 8 and 16 months of age, we next investigated the swimming performance at 12 months using a swim tunnel assay (Loligo® Systems Swim Mini). After placing the fish in the swim tunnel as shown in Fig. 4K, we allowed 20 minutes for habituation. A water flow current was initiated within the swim tunnel and gradually increased using reproducible preset speed settings. The starting speed was set at 1 and then gradually increased to 1.5, 2, 2.5, 3, 3.5, 4, 4.5, and 5, every 2 minutes. The experiment was recorded and analyzed to determine the time spent swimming against the water current until the fish gave up and hit the baffle located at the end of the swim tunnel three times (Fig. 4L). Using 8 males and 8 females per group (n = 16 per group) at 12 months of age, we found that all homozygous KO fish showed lower swimming performance than WT fish, with *atxn1a* KO fish showing the lowest performance (Fig. 4M). Reasons for poor swimming performance could be related to deficits in strength, endurance, coordination and/or motivation.

### RNA-seq reveals shared and gene-specific transcriptomic alterations across KO larvae

We applied RNA-seq to broadly investigate whether KO of *atxn1a*, *atxn1b*, and *atxn1l* genes results in transcriptional alterations at 5 dpf. At this time point, the basic organization of the central nervous system is complete in zebrafish, and all *atxn1* paralogs are expressed in the brain, as previously shown by *in situ* hybridization (Vauti et al., 2021). Twenty larvae per group were pooled for RNA extraction and treated as a single sample (Fig. 5A). This process was repeated four times on separate days, with samples collected at the same time of day to minimize circadian rhythm effects. This resulted in four samples per group. After checking RNA quality and quantity, paired-end sequencing was performed, and the data were analyzed on the Galaxy platform for quality control, alignment, read counting, and identification of differentially expressed genes (DEGs) (The Galaxy Community et al., 2024).

**Fig. 5.**
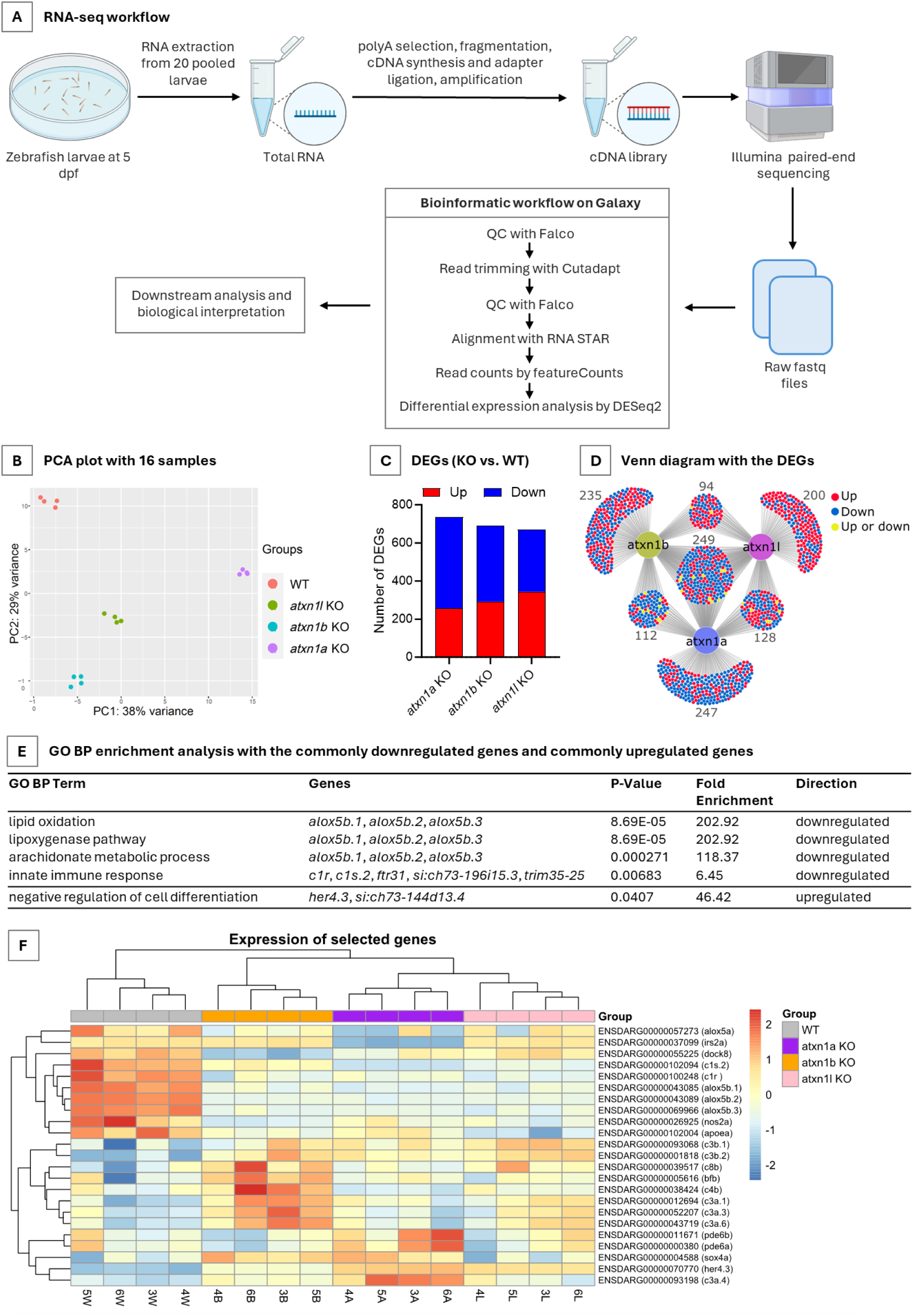
Transcriptomic analysis of 5 dpf larvae. **A.** Sample collection, library preparation, sequencing, and bioinformatic analysis workflow for transcriptomic analysis using RNA-seq. Each sample was obtained from 20 pooled larvae at 5 dpf. A total of 16 samples (4 per group) were collected. **B.** Principal component analysis (PCA) plot of the 16 RNA-seq samples. **C.** Number of significant differentially expressed genes (DEGs; adjusted P value < 0.05 and |log2FC| > 1) in WT vs. *atxn1a* KO, WT vs. *atxn1b* KO, and WT vs. *atxn1l* KO comparisons. **D.** Venn diagram showing common and uniquely differentially expressed genes across the different KO groups. **E.** Gene Ontology Biological Process (GO BP) enrichment analysis with the commonly downregulated and commonly upregulated genes across the three KO groups. **F.** Heatmap showing the expression of selected genes.

Principal component analysis (PCA) showed clear clustering of samples according to groups (Fig. 5B). The first two components, PC1 and PC2, explained 38% and 29% of the variability in the expression data (Supplementary Table 1), respectively. Differential expression analysis was performed for each KO line vs. WT (Supplementary Table 2). Using the criteria of FDR value < 0.05 and |log2FC| > 1 (Supplementary Fig. 4-6), we identified 736 (256 upregulated, 480 downregulated) DEGs in *atxn1a* KO, 690 (292 upregulated, 398 downregulated) DEGs in *atxn1b* KO, and 671 (344 upregulated, 327 downregulated) DEGs in *atxn1l* KO samples (Fig. 5C). Overall, we identified 1,265 DEGs when the DEGs from these three analyses were combined. Among these, 249 DEGs were common to all three KO groups (Fig. 5D). Of these 249 genes, 238 (238/1,265 = 18.9%) DEGs showed the same direction of change, with 157 downregulated and 81 upregulated genes. Alongside these shared DEGs, we identified 247 (19.5%), 235 (18.6%), and 200 (15.8%) DEGs that were unique to *atxn1a*, *atxn1b*, and *atxn1l* KO respectively. These findings indicate that KO of each of these genes results in a markedly altered transcriptome, with both shared and KO-specific changes.

### Shared and uniquely enriched biological processes revealed by Gene Ontology enrichment analysis

In order to make biological sense of the DEGs uncovered, we performed Gene Ontology (GO) Biological Process (BP) enrichment analysis using the commonly downregulated and commonly upregulated genes (FDR value < 0.05 and |log2FC| > 1) across *atxn1a*, *atxn1b*, and *atxn1l* KOs. This enrichment analysis was performed with the DAVID web server (Huang et al., 2009; Sherman et al., 2022). The commonly downregulated genes showed significant enrichment of GO BP terms related to the lipoxygenase pathway (lipid oxidation (GO:0034440), lipoxygenase pathway (GO:0019372), arachidonate metabolic process (GO:0019369)) (Fig. 5E). This enrichment was driven by multiple *alox5* genes (Fig. 5F), which encode arachidonate 5-lipoxygenase, a key enzyme in leukotriene biosynthesis (Sun et al., 2019), and is expressed in Purkinje cells (Chen et al., 2010). In addition, the GO BP term “innate immune response” (GO:0045087), which includes classical complement pathway component genes such as *c1r* and *c1s.2* (Fig. 5F), was also enriched among the commonly downregulated genes. In contrast, the commonly upregulated genes were enriched for the GO BP term “negative regulation of cell differentiation” (GO:0045596), driven by increased expression of *her4.3* (Fig. 5F). *her4.3* is a zebrafish ortholog of *Hes5* (Wang et al., 2024); studies of HES5 in mice have shown that it inhibits neuronal differentiation and suppresses proliferation of neural progenitors (Bansod et al., 2017). Collectively, these findings suggest a possible suppression of leukotriene metabolism and enhanced repression of cellular differentiation programs across the KO groups.

Next, we performed GO BP enrichment analysis using the danRerLib Python package (Schwartz et al., 2024). The package allows the application of a logistic regression–based approach to analyze the full differential expression results rather than a preselected list of DEGs and detects significantly upregulated and downregulated pathways. The top 5 downregulated GO BP terms in the *atxn1a* KO, *atxn1b* KO, and *atxn1l* KO are shown in Fig. 6A, 6B, and 6C, respectively (full list of enriched pathways are provided as Supplementary Table 3). We found that “dendritic cell migration” (GO:0036336) and “regulation of cytokine production involved in inflammatory response” (GO:1900015) were commonly enriched in all three KOs. These two GO BP terms include *alox5* family genes, which, as mentioned previously, were commonly downregulated in all KOs. *dock8*, which belongs to GO:0036336, was also downregulated, particularly in the *atxn1a* and *atxn1b* KOs (Fig. 5F). Loss of *dock8* has been reported to attenuate microglial colonization in early zebrafish larvae (Wu et al., 2022). In addition to *alox5* family genes, GO:1900015 also includes *nos2a*. *nos2a*, an ortholog of human *NOS2* encoding inducible nitric oxide synthase, was commonly downregulated in all KOs (Fig. 5F). This finding is consistent with reduced levels of inducible nitric oxide synthase in an experimental autoimmune encephalomyelitis model in hemizygous *Atxn1* (*Atxn1*^2Q/−^) mice (Talukdar et al., 2025). Together, these shared findings suggest potential suppression of leukotriene and nitric oxide biosynthesis, as well as delayed migration of dendritic cells/ macrophages/ microglia.

**Fig. 6.**
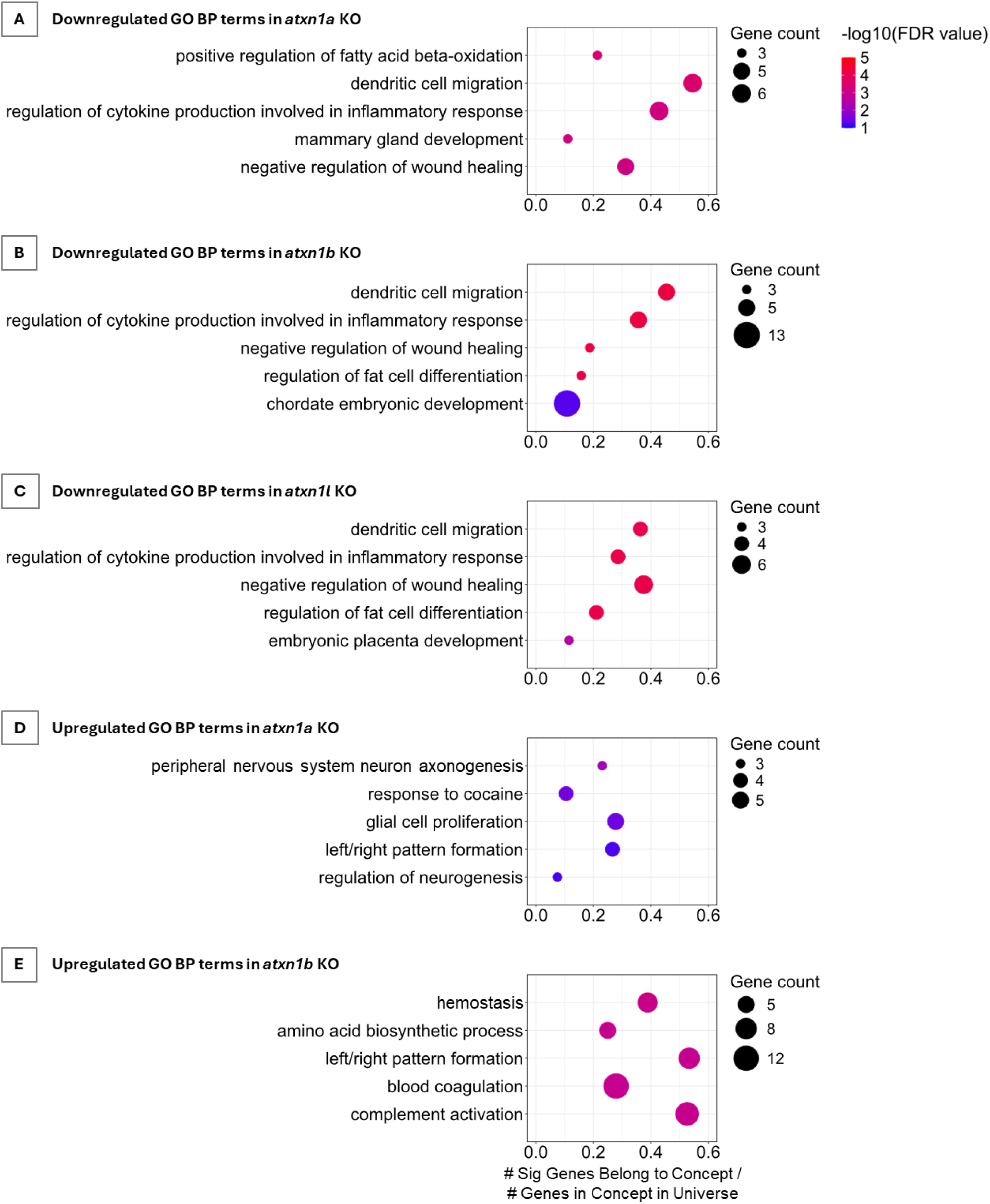
Downregulated and upregulated GO BP terms in KO groups. **A–C.** Downregulated Gene Ontology Biological Process (GO BP) terms in *atxn1a* KO (**A**), *atxn1b* KO (**B**), and *atxn1l* KO (**C**) larvae. Only top 5 terms with at least 3 genes are shown (full list is provided in the Supplementary Table 3). **D, E.** Upregulated GO BP terms in *atxn1a* KO (D) and *atxn1b* KO (**E**) larvae. Only top 5 terms with at least 3 genes are shown (full list is provided in the Supplementary Table 3).

We next explored beyond shared downregulated processes to identify GO BP uniquely associated with each KO line. Interestingly, *atxn1a* KO showed unique downregulation of “positive regulation of fatty acid beta-oxidation” (GO:0032000). This term includes *irs2a*, which was significantly downregulated in the *atxn1a* KO (log2FC = −4.8, FDR = 0), but not in the *atxn1b* or *atxn1l* KOs (Fig. 5F). This finding may indicate altered insulin signaling and impaired brain development, as insulin signaling has been reported to promote neurogenesis in zebrafish (Gence et al., 2023), whereas knockout of *Irs2* in mice results in impaired brain growth (Schubert et al., 2003).

The top 5 upregulated GO BP terms in the *atxn1a* KO and *atxn1b* KO from the enrichment analysis using danRerLib are shown in Fig. 6D and 6E. *atxn1l* KO did not show upregulated GO BP terms with FDR value <0.1 (Supplementary Table 3). We observed that the *atxn1a* and *atxn1b* KOs showed significant enrichment of multiple GO BP terms, most of which were unique to each KO line. Several nervous system–related terms were enriched in the *atxn1a* KO, including “peripheral nervous system axonogenesis” (GO:0048936) and “glial cell proliferation” (GO:0014009). GO:0014009 includes *sox4a*, which was significantly upregulated in both *atxn1a* and *atxn1b* KOs (Fig. 5F). Previous studies have shown that SOX4, the human ortholog of sox4a, represses oligodendrocyte differentiation from neural progenitors (Braccioli et al., 2018), whereas in mice, prolonged SOX4 expression causes Bergmann glia to fail to establish radial glial morphology (Hoser et al., 2007). The *atxn1b* KO showed enrichment of “complement activation” (GO:0006956). This term includes orthologs of complement pathway component genes *c3a*, *c3b*, *c4b*, and *c8b* which were significantly upregulated in *atxn1b* KO (Fig. 5F).

### WGCNA analysis reveals highly correlated gene modules specific to individual *atxn1* gene KOs

To further interrogate the biological meaning of the RNA-seq data, we used the CEMiTool to perform weighted gene co-expression network analysis (WGCNA) (Cheng et al., 2020; Russo et al., 2018). WGCNA identifies groups of genes, called modules, which have correlated expressions and are likely to share biological functions or regulatory mechanisms. This analysis identified 12 modules, designated M01 to M12 (Supplementary Table 4, Supplementary Fig. 7-18). Module–trait relationship analysis revealed several strongly correlated modules, including M01 with WT, M03 with *atxn1a* KO, M07 with *atxn1b* KO, and M08 with *atxn1l* KO (Fig. 7A).

**Fig. 7.**
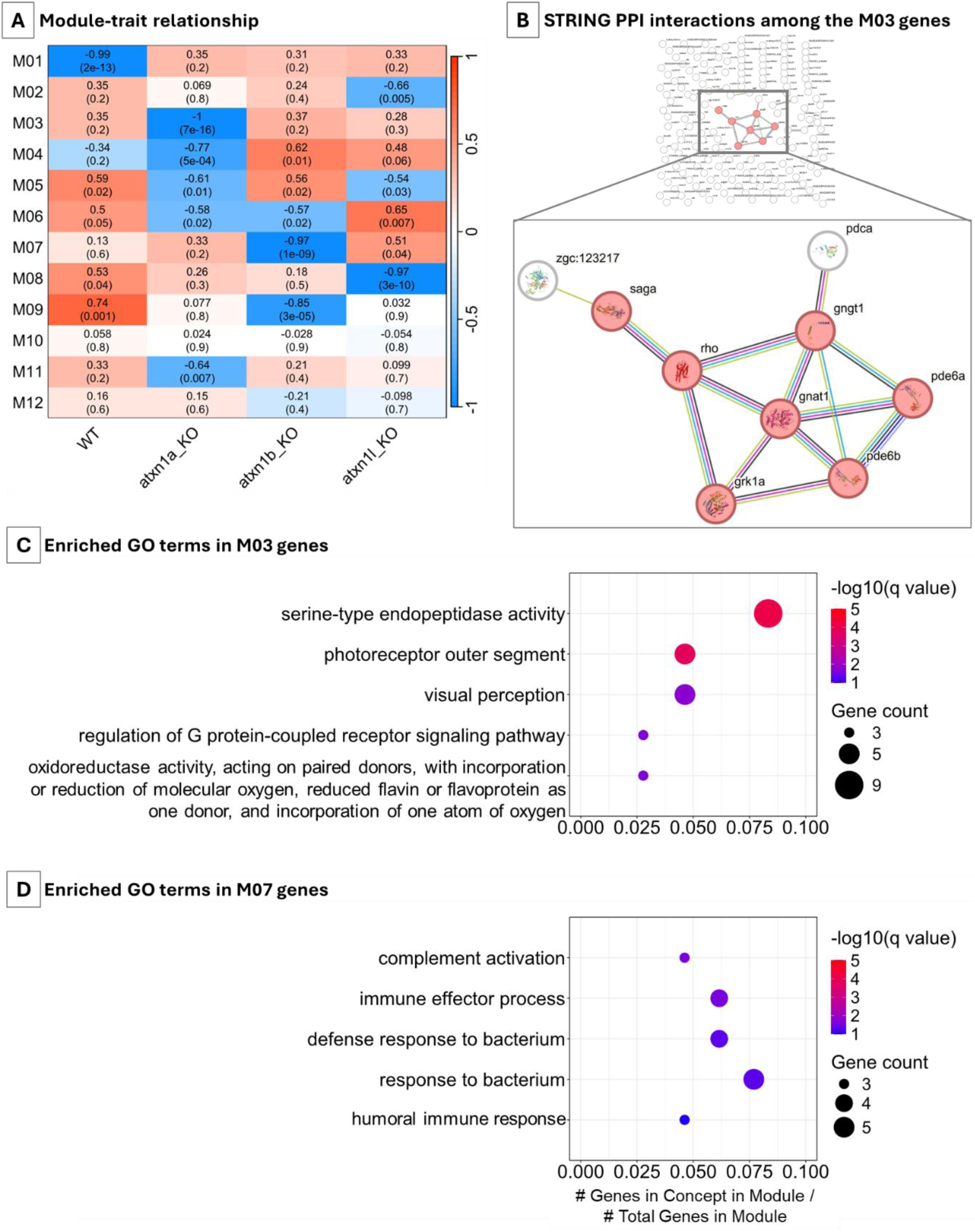
Weighted gene co-expression network analysis (WGCNA). **A.** Module–trait relationships determined from the eigengene values of the 12 modules (M01–M12) detected by CEMiTool. **B.** STRING protein-protein interaction (PPI) analysis with the M03 genes (PPI enrichment p-value = 1.97e-06; minimum interaction score = 0.7). Proteins related to KEGG pathway “Phototransduction” (dre04744) are highlighted in reddish color. **C, D.** Top 5 overrepresented GO terms in M03 (**C**) and M07 (**D**). Full list is provided in the Supplementary Table 5.

We assessed the gene list in each module for potential protein–protein interaction (PPI) with STRING (Szklarczyk et al., 2023) and found an interaction cluster in the M03 module (Fig. 7B; PPI enrichment p-value = 1.97e-06; minimum interaction score = 0.7). Most of these genes belonged to the KEGG pathway “Phototransduction” (dre04744; FDR = 5.75e-06).

We also performed over-representation analysis using CEMiTool to identify significantly enriched GO terms within each module (Supplementary Table 5). Module M03 was enriched for GO terms related to photoreceptors and visual perception (Fig. 7C). This finding is in line with the light-dependent locomotor phenotype described earlier for *atxn1a* KO larvae. Notably, this module includes *pde6a* and *pde6b*, which showed higher expression in the *atxn1a* KO (Fig. 5F). These genes encode components of key phosphodiesterase enzyme in the phototransduction cascade (Cote, 2021; Muradov et al., 2010).

Module M07 showed enrichment for complement pathway–related GO terms (Fig. 7D) and included *bfb* (orthologous to human *CFB*), which showed higher expression in the *atxn1b* KO (Fig 5F). Module M08 was enriched for lipoprotein-related GO terms (Supplementary Table 5), which included the *APOE* ortholog *apoea*. *apoea* had lower expression in all KOs compared to the WT, with the lowest expression observed in the *atxn1l* KO (Fig 5F).

## Discussion

Over three decades have passed since the discovery that a CAG repeat expansion in *ATXN1* causes SCA1 (Orr et al., 1993). While significant progress has been made in elucidating its pathogenesis, a disease-modifying therapy for SCA1 remains an unmet clinical need. Moreover, emerging evidence has expanded the clinical relevance of *ATXN1* beyond SCA1, linking it to Alzheimer’s disease (Suh et al., 2019), amyotrophic lateral sclerosis (Tazelaar et al., 2020), and multiple sclerosis (Carver et al., 2025; Didonna et al., 2020; Ma & Didonna, 2023). Current preclinical therapeutic strategies for SCA1 largely focus on reducing ATXN1 levels, both mutant and wild-type, through approaches such as antisense oligonucleotides or RNA interference (Kerkhof et al., 2023). Inhibition of ATXN1 S776 phosphorylation and ATXN1L overexpression have also been examined (Kerkhof et al., 2023; Srinivasan & Shakkottai, 2019).

While various mouse models expressing polyQ-expanded ATXN1 have provided invaluable insights into toxic gain-of-function mechanisms, the physiological role of the WT protein remains comparatively understudied. *Atxn1* KO mice exhibit cognitive and motor deficits and an altered cerebellar proteome (Matilla et al., 1998; Sánchez et al., 2016). *ATXN1* loss of function also potentiates Alzheimer’s disease (Suh et al., 2019) and multiple sclerosis pathogenesis (Didonna et al., 2020; Talukdar et al., 2025). These findings, together with the fact that some proposed therapeutic strategies may interfere with the WT copy of ATXN1 (Kerkhof et al., 2023), underscore the need to study ATXN1 function, expand the repertoire of KO models beyond mice, and investigate the biological consequences of its depletion. Zebrafish offer advantages as an alternative or complementary model to mice due to their rapid embryonic development, transparent embryos that enable advanced live imaging, high-throughput behavioral assessment at both larval and adult stages, and suitability for high-throughput drug screening (Sarasamma et al., 2023). To date, the zebrafish *atxn1*-related repertoire has been limited to a transgenic Purkinje cell–specific model expressing patient-derived Atx1[82Q] (Elsaey et al., 2021). While this model is useful for studying polyQ toxicity, the field lacks KO zebrafish models to investigate the physiological functions of *ATXN1* and *ATXN1L* orthologs.

In this study, we used CRISPR–Cas9 genome editing to introduce frameshift mutations in *atxn1a*, *atxn1b*, and *atxn1l* genes and established three different lines for characterization. After establishing the mutant lines, we assessed survival, hatching rate, and body length up to 5 dpf. We observed that all KO groups had lower survival rates due to increased mortality by 1 dpf. However, from 1 to 5 dpf, there was minimal death. This increased mortality could be due to lower-quality germ cells produced by the KO fish compared to WT, or to early developmental defects occurring in the initial hours after fertilization. In our study, we found that nearly all embryos hatched by 72 dpf in all groups which is consistent with previous reports of hatching at 2-3 dpf (Wisenden et al., 2022). However, at 2 dpf, we observed a relatively lower hatching rate in *atxn1a* KO embryos compared to WT.

Alternating light–dark cycles are a commonly used method to assess locomotor behavior in zebrafish larvae (Tuz-Sasik et al., 2022). The duration of the light and dark phases varies across studies. In the current study, we designed two paradigms: a short paradigm in which each light or dark phase was 10 minutes, and a long paradigm in which each phase was 1 hour. While the short paradigm failed to produce any reproducible phenotypic differences, the long paradigm showed a light-dependent decrease in the distance traveled by *atxn1a* KO larvae, whereas *atxn1b* and *atxn1l* KOs showed a global reduction spanning both light and dark conditions. In line with this, RNA-seq data revealed an *atxn1a* KO-associated gene module enriched for phototransduction and vision-related genes. This module includes the *pde6a* and *pde6b*, which showed relatively higher expression in *atxn1a* KO larvae. These genes encode components of a key photoreceptor phosphodiesterase enzyme responsible for regulating the phototransduction cascade (Cote, 2021; Muradov et al., 2010). Under dark conditions, high cGMP levels maintain photoreceptors in a depolarized state and promote glutamate release. Upon exposure to light, the phototransduction cascade is initiated, resulting in activation of phosphodiesterase, which hydrolyzes cGMP. Reduced cGMP levels hyperpolarize photoreceptors, decrease glutamate release, and enable light perception (Iribarne & Masai, 2017). Therefore, it is tempting to hypothesize that the increased phosphodiesterase levels observed in *atxn1a* KO larvae reduce cGMP levels in photoreceptors. This may lead to amplified light perception and reduced movement. Further studies are required to test this hypothesis.

We assessed adult fish behavior using the novel tank test and swim tunnel assay. Consistent with previous reports (Golushko et al., 2025), we found that WT adult fish showed a tendency to occupy the bottom of the tank. Confinement to the bottom zone is a natural anxiety-like behavior, whereas exploration of the top of the tank is considered an indicator of reduced anxiety (Golushko et al., 2025). Interestingly, *atxn1a* KO fish showed a pronounced increase in time spent at the top of the tank, consistent with a reduced anxiety-like behavior. This is in line with *Atxn1* null mice showing cognitive deficits such as impaired learning and reduced anxiety (Lee et al., 2011; Matilla et al., 1998). This finding may also reflect cognitive impairment, as RNA-seq analysis revealed dysregulation of gene expression that may impact brain development and function (discussed below). Furthermore, this behavioral deficit demonstrates a gradient of phenotypic severity with *atxn1a* KOs displaying the earliest and most pronounced alteration in vertical tank exploration, followed by *atxn1b* and then *atxn1l* mutants; with each line showing progression in dysfunction over time. In the swim tunnel assay, we observed that all KO groups exhibited reduced swimming performance, with *atxn1a* KO showing the lowest performance. This reduction in swimming performance may be attributable to impairments in muscular strength, endurance, motor coordination, and/or motivational drive.

Transcriptomic analysis suggests an important role of the *atxn1*-family genes in neurodevelopment and neurodegeneration risk. We observed robust downregulation of *irs2a* (insulin receptor substrate 2a) in *atxn1a* KO larvae. Insulin and insulin-like growth factors mediate phosphorylation of insulin receptor substrates upon binding to insulin receptors. Previous study in mice showed that IRS2 signaling is important for neuronal proliferation and increasing brain size during development (Schubert et al., 2003). Although studies of *Irs2* orthologs in zebrafish are scarce, it was shown that insulin signaling promotes neurogenesis in zebrafish (Gence et al., 2023). Therefore, it will be interesting to further investigate the consequences of *irs2a* downregulation in zebrafish neurodevelopment. Another interesting transcriptomic change is the downregulation of *dock8*. This was seen in all KO groups, but particularly pronounced in the *atxn1a* and *atxn1b* KOs. A previous study showed that *dock8* deficiency attenuates microglial colonization in early zebrafish larvae (Wu et al., 2022). Microglia play a crucial role in brain development by regulating synaptic pruning, engulfing apoptotic neurons, and modulating neuronal activity. Therefore, we hypothesize that the downregulation of *dock8* may result in impaired microglial colonization of the brain. Further research is necessary to explore this hypothesis and to investigate whether it results in impaired brain development and/or increases the risk of neurodegeneration. Finally, *nos2a*, a gene encoding inducible nitric oxide synthase, was downregulated in all KOs. NOS2 is known to play regulatory roles in immunity and neurodegenerative diseases (Bogdan, 2015; Iova et al., 2023; Sonar & Lal, 2019). Genetic deletion of *NOS2* in Alzheimer’s models leads to increased insoluble Aβ, tau hyperphosphorylation, neuroinflammation, and enhanced neuronal death (Colton et al., 2006, 2008; Kim et al., 2023).

In summary, we generated the first *atxn1a*, *atxn1b*, and *atxn1l* knockout zebrafish models and characterized their phenotypic and transcriptomic alterations. We found that *atxn1a* KO larvae exhibit a specific light-dependent reduction in locomotor activity in a long light–dark paradigm. Consistent with this phenotype, the gene expression profile for *atxn1a* KO larvae is highly correlated with a gene module enriched for genes regulating photoreceptor and visual function. Transcriptomic analysis revealed both shared and KO-specific alterations among the three KO lines. Although the RNA-seq samples were derived from whole embryos, many of the DEGs and enriched biological processes were related to the nervous system, indicating an important role for this family of genes in nervous system development and function. Dysfunction in pathways regulating neuronal proliferation, microglial migration and neuroinflammation seen in this study may shed new light in existing research connecting loss of *ATXN1* function to increased Alzheimer’s disease and multiple sclerosis risk. Furthermore, better understanding the functional consequences of reduced *ATXN1* expression will be paramount in developing safe and effective therapeutic strategies for SCA1.

## Methods

### Zebrafish

Zebrafish were maintained at 28.5 °C under a 14-h light/10-h dark cycle at the Baylor College of Medicine (BCM) Zebrafish Research Facility, using a recirculating water system (Tecniplast USA). WT zebrafish were of the AB strain (Westerfield, 2000). The *atxn1a* mutant lines were generated in the AB background at BCM. The *atxn1b* and *atxn1l* mutant lines were generated in the AB background at the University of Utah Health Science Core Facilities and the embryos were subsequently transferred to BCM. All experimental procedures were approved by the BCM Institutional Animal Care and Use Committee.

### Embryo collection

Adult zebrafish were allowed to spawn naturally either as pairs or in groups. Embryos were collected at 20-minute intervals to maintain developmental synchrony within each cohort and between groups. Collected embryos were transferred to 60 cm² Petri dishes at a density not exceeding 100 embryos per dish, maintained in 0.5× E2B medium with 0.5 mg/L methylene blue, and incubated at 28.5 °C.

### Generation of mutant zebrafish using CRISPR/Cas9

sgRNAs targeting the first coding exons of *atxn1a*, *atxn1b*, and *atxn1l* genes were designed using CHOPCHOP (Montague et al., 2014) and purchased from Integrated DNA Technologies (IDT). Glass needles for microinjection were prepared with a Sutter Instruments Fleming/Brown Micropipette Puller (model P-97) and a regulated air-pressure microinjector (Harvard Apparatus, PL1–90). Embryos were injected at the 1-cell stage into the cytoplasm with Cas9 protein (EnGen® Spy Cas9 NLS, cat. #M0646T) and sgRNA (800 ng/µL Cas9, 0.05% phenol red, and 322.44 [*atxn1a*], 322.44 [*atxn1b*], or 323.46 [*atxn1l*] ng/µL sgRNA). Injected embryos (F_0_) were raised to adulthood and crossed with WT AB fish to generate F_1_ embryos which were raised. F_1_ offsprings were sequenced by tail fin biopsies to identify frameshift mutations predicted to cause loss of protein function.

### Genomic DNA isolation

500 µL of lysis buffer (50 mM Tris-HCl [pH 8.0], 100 mM EDTA, 100 mM NaCl, and 1% SDS in Milli-Q water) and 25 µL of 10 mg/mL proteinase K (GoldBio, cat. #P-480-1) were added to embryo or tail biopsy samples. After vortexing, samples were incubated overnight at 55 °C with shaking. The samples were then centrifuged for 5 minutes at 11,800 rpm at room temperature. Next, 400 µL of the supernatant was transferred to new tubes containing 200 µL of salt solution (4.21 M NaCl, 0.63 M KCl, and 10 mM Tris-HCl [pH 8.0] in Milli-Q water). After vortexing, samples were centrifuged for 5 minutes at 11,800 rpm. Then, 400 µL of the supernatant was transferred to new tubes containing 600 µL of 100% ethanol. After vortexing, the samples were centrifuged and the supernatants were discarded. Subsequently, 500 µL of 80% ethanol was added, followed by vortexing and centrifugation for 5 minutes at 11,800 rpm. The supernatants were discarded, and the samples were allowed to air-dry for 20 minutes. Finally, 100 µL of Milli-Q water was added to the samples, which were then stored at −20 °C until further use.

### PCR

PCR reaction mixes with a total volume of 24 µL were prepared using 12 µL EconoTaq® PLUS GREEN 2X Master Mix (cat. #30033-1), 2 µL of 10 µM primer mix, 2 µL of template DNA solution, and Milli-Q water to the final volume. PCR was performed in a Bio-Rad T100™ Thermal Cycler. The PCR protocol consisted of 95 °C for 3 minutes, followed by 33 cycles of 95 °C for 30 seconds, 60 °C for 40 seconds, and 72 °C for 2 minutes, with a final extension at 72 °C for 5 minutes. PCR products were analyzed by 1% agarose (Fisher BioReagents™, cat. #BP160-500) gel electrophoresis.

### Sanger sequencing

The mutation at each gene’s target site was confirmed by Sanger sequencing. PCR products were purified using NucleoSpin® PCR Clean-up (Takara Bio, cat. #740609.250) following the manufacturer’s instructions. Then, 5 µL of purified PCR product (30 ng/µL) was mixed with 5 µL of 10 µM primer (forward or reverse) and submitted to Eurofins Genomics for Sanger sequencing.

### Embryo and larval phenotyping

For each experiment, spawning was performed for WT, *atxn1a*, *atxn1b*, and *atxn1l* KO lines. Viable embryos were collected at 2 hours post-fertilization and placed in Petri dishes (50 embryos per dish). Each experiment started with 100 embryos and was followed until 5 dpf to assess survival and hatching. The number of live embryos was recorded at 1, 2, 3, and 5 dpf. At the same time, the number of hatched embryos was also recorded. This experimental procedure was repeated 10 times. For the first four experiments, images of 10 embryos per group were taken, one at a time, at 5 dpf to measure larval body size. Imaging was done with ECHO RVL-100-M microscope. Body length from head to tail was measured with ImageJ.

### Light-dark challenge assay

At 5 dpf, randomly selected larvae were placed individually into wells of a flat-bottom 96-square-well plate (Whatman™, cat. #7701-1651). Each well contained 225 µL of 0.5× E2B medium with 0.5 mg/L methylene blue. In each experiment, 48 WT larvae and 48 KO larvae from a given line were used. The plate was then placed in a DanioVision observation chamber (Noldus, Wageningen, Netherlands), equipped with infrared illumination from below and an infrared-sensitive camera positioned above the plate. Light–dark paradigm was then initiated, consisting of a habituation period followed by alternating light and dark phases (Fig. 3A). Light intensity level during the light phase was set to 100%, and the temperature was maintained at 28.5 °C. A total of five experiments with larvae of different cohorts were conducted for each KO line. All experiments were initiated at the same time of day, between 12:00 pm and 1:00 pm.

The distance traveled by each larva per second was calculated from the recorded videos using EthoVision XT software (Version 13; Noldus, Wageningen, Netherlands). The data were further analyzed in the R environment (Version 4.2.3) to calculate the distance traveled over each 2-minute interval as well as the distance traveled during each phase. These data were then exported from R and visualized using GraphPad Prism (Version 10). Statistical comparisons between WT and KO larvae were also performed using GraphPad Prism.

### Novel tank test

Adult zebrafish were subjected to a novel tank diving assay following a recently published protocol (Keerthisinghe et al., 2026). WT, *atxn1a*, *atxn1b*, and *atxn1l* KO fish were tested at 8, 12, and 16 months of age. Individual fish were placed in a novel tank (20 cm L × 12 cm H × 9 cm W; ≈1.8 L) filled with fresh system water. After 5 minutes of habituation, video recording was started and continued for 10 minutes. Videos were acquired using an iPhone 15 camera positioned 20 cm away from the tank. Recordings were analyzed using a previously trained DeepLabCut pose estimation model (DeepLabCut v3.0.0-rc6) as described (Keerthisinghe et al., 2026). Positional tracking data were extracted, and the tank was vertically divided into three equal zones (bottom, middle, and top). The duration spent in each zone was quantified to determine region occupancy. Representative heatmaps were generated from the tracking data using a Python script (available at https://zenodo.org/records/17624893).

### Swim tunnel assay

Swimming performance was assessed using a swim tunnel respirometer (Swim Tunnel Mini, Loligo® System). WT, *atxn1a*, *atxn1b*, and *atxn1l* KO zebrafish were tested at 12 mpf. For each genotype, sixteen fish (8 males and 8 females) were analyzed. Fish were placed individually in the swim chamber containing system water from the zebrafish facility. Water temperature was maintained at 28.5 °C using an integrated heater. Fish were allowed to acclimate in the chamber for 20 minutes. Following acclimation, water velocity was initially set at a low speed (speed 1) and then increased in a stepwise manner by manually adjusting the propeller speed every 2 minutes. At each interval, speed was incrementally increased by 0.5 units on the controller dial. Fish were required to swim continuously against the current, and performance was determined as the time elapsed until a predefined fatigue criterion was reached, defined as hitting the rear baffle of the swim chamber with its tail three times.

### RNA extraction

A total of 20 larvae (5 dpf) from each group were taken randomly from the Petri dish into a 1.5 mL microcentrifuge tube and placed on ice. This pool of 20 larvae was considered one sample. In each batch, we collected one sample from each group, and the process was repeated four times on separate days to collect four samples per group. After pooling the 20 larvae, 1 mL of TRIzol™ reagent (Invitrogen, cat. #15596018) was added to the tube. Larvae were ground by pipetting up and down with a syringe attached to a 20G needle (BD PrecisionGlide™) 20 times and then 10 times with a 26G (BD PrecisionGlide™) needle. The samples were then incubated for 5 minutes in TRIzol at room temperature. Then, 200 µL chloroform (Sigma-Aldrich, cat. #319988-500ML) was added to the sample tubes. The tubes were shaken gently for 15 seconds by hand and then incubated for 2 minutes at room temperature. Next, centrifugation was done for 30 minutes at 13,000 rpm at 4 °C. A 200 µL volume from the upper clear phase from each tube was transferred to new tubes containing 500 µL isopropanol (Sigma-Aldrich, cat. #675431-4L) and stored at -20 °C overnight. The next day, samples were allowed to thaw and then centrifuged for 15 minutes at 13,000 rpm at 4 °C. The supernatant was removed, and the pellet was washed with 1 mL of pre-chilled 75% ethanol (Koptec, cat. #V1016). After gentle shaking, samples were centrifuged for 10 minutes at 10,000 rpm at 4 °C. The supernatant was removed, and the samples were allowed to dry for 10–20 minutes. Then, 100 µL of nuclease-free water (Invitrogen, cat. #AM9938) was added and pipetted up and down 20 times to dissolve the RNA pellet. RNA concentration was measured using NanoDrop™ One (ThermoFisher), and the samples were stored at -80 °C until further use.

### RNA-seq data analysis

Raw RNA-seq data (FASTQ files) were analyzed on Galaxy (Version 1.2.4+galaxy0) for quality control, alignment, read counting, and identification of DEGs (The Galaxy Community et al., 2024). Briefly, the quality of the FASTQ files was checked using Falco (Version 1.2.4) (De Sena Brandine & Smith, 2021). Adapter trimming and removal of low-quality reads were performed with Cutadapt (Version 5.1) (Martin, 2011). Reads were aligned to the zebrafish reference genome (GRCz11.dna.primary_assembly.fa.gz) using RNA STAR (Version 2.7.11b) (Dobin et al., 2013). Gene expression was quantified from the alignment output files (BAM files) using featureCounts (Version 2.1.1) (Liao et al., 2014). Finally, differential expression analysis was performed with DESeq2 (Version 2.11.40.8) (Love et al., 2014).

### Enrichment analysis

Enrichment analysis was done separately for different steps in the RNA-seq data analysis workflow, as appropriate. With the criteria of FDR < 0.05 and |log2FC| > 1, we identified significantly upregulated and downregulated genes in all WT vs KO comparisons. Then, we identified the commonly upregulated and commonly downregulated genes across these lists. Enrichment analysis for GO BP terms in these commonly upregulated or downregulated genes was performed using the DAVID web server (Huang et al., 2009; Sherman et al., 2022).

Enrichment analysis for GO BP terms using each of the differential expression (DE) analysis results was performed with danRerLib (Schwartz et al., 2024). A logistic regression model and the “variable” option for “org” were chosen for this analysis. The “variable” option uses true zebrafish ontology annotations for genes where available and maps genes to human orthologs for annotation only if zebrafish annotation does not exist for that gene. The analysis was done separately for the DE results of *atxn1a*, *atxn1b*, and *atxn1l*. The top 5 upregulated and top 5 downregulated pathways were plotted in R after filtering out terms containing fewer than 3 genes.

### WGCNA

WGCNA analysis was performed using webCEMiTool (Russo et al., 2018) with expression data from all 16 samples. Genes with fewer than 10 reads in more than 50% of samples were filtered out. Then, the VST-normalized data of the remaining genes from the DESeq2 output were used as input to CEMiTool. A GO and pathway database was also provided to CEMiTool for enrichment analysis of the detected co-expression gene modules. This database was obtained from g:Profiler (Raudvere et al., 2019) and contained GO, KEGG, Reactome, and WikiPathways data. Heatmaps of the module genes were created in the R environment using the pheatmap (Version 1.0.13) package.

## Abbreviations

atxn1: ataxin 1
atxn1l: ataxin 1 like
SCA1: spinocerebellar ataxia 1
KO: knockout
dpf: days post-fertilization
WT: wild-type
polyQ: polyglutamine

## CRediT authorship contribution statement

Anwarul Karim: Conceptualization; Methodology; Software; Validation; Formal analysis; Investigation; Visualization; Writing - Original Draft

Pramuk Keerthisinghe: Methodology; Software; Formal analysis; Investigation; Visualization; Writing - Original Draft

Sreeja Sarasamma: Methodology; Investigation

Nicholas A. Ciaburri: Investigation

Mabel Guerra Giraldez: Investigation

Kylan Naidoo: Investigation

James P. Orengo: Conceptualization; Methodology; Resources; Writing - Review & Editing; Supervision; Project administration; Funding acquisition

All the authors approved the final version of the manuscript.

---

**Supplementary Fig. 1.**
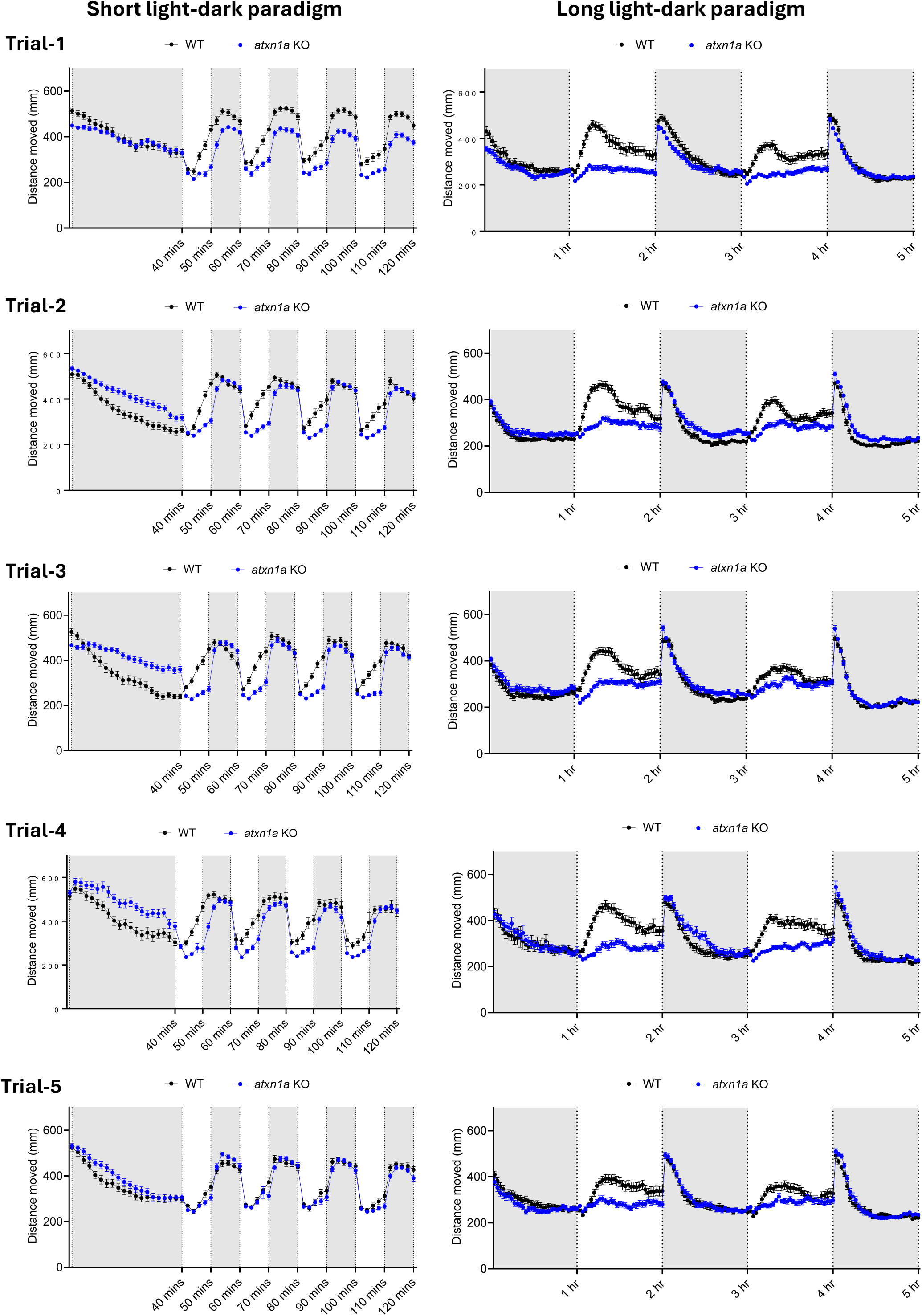
Assessment of locomotor activity of homozygous *atxn1a* KO larvae at 5 dpf. All five trials are shown. Left panel represents the short light-dark paradigm and the right panel represent long light-dark paradigm. Each trial was performed with 48 WT and 48 KO larvae.

**Supplementary Fig. 2.**
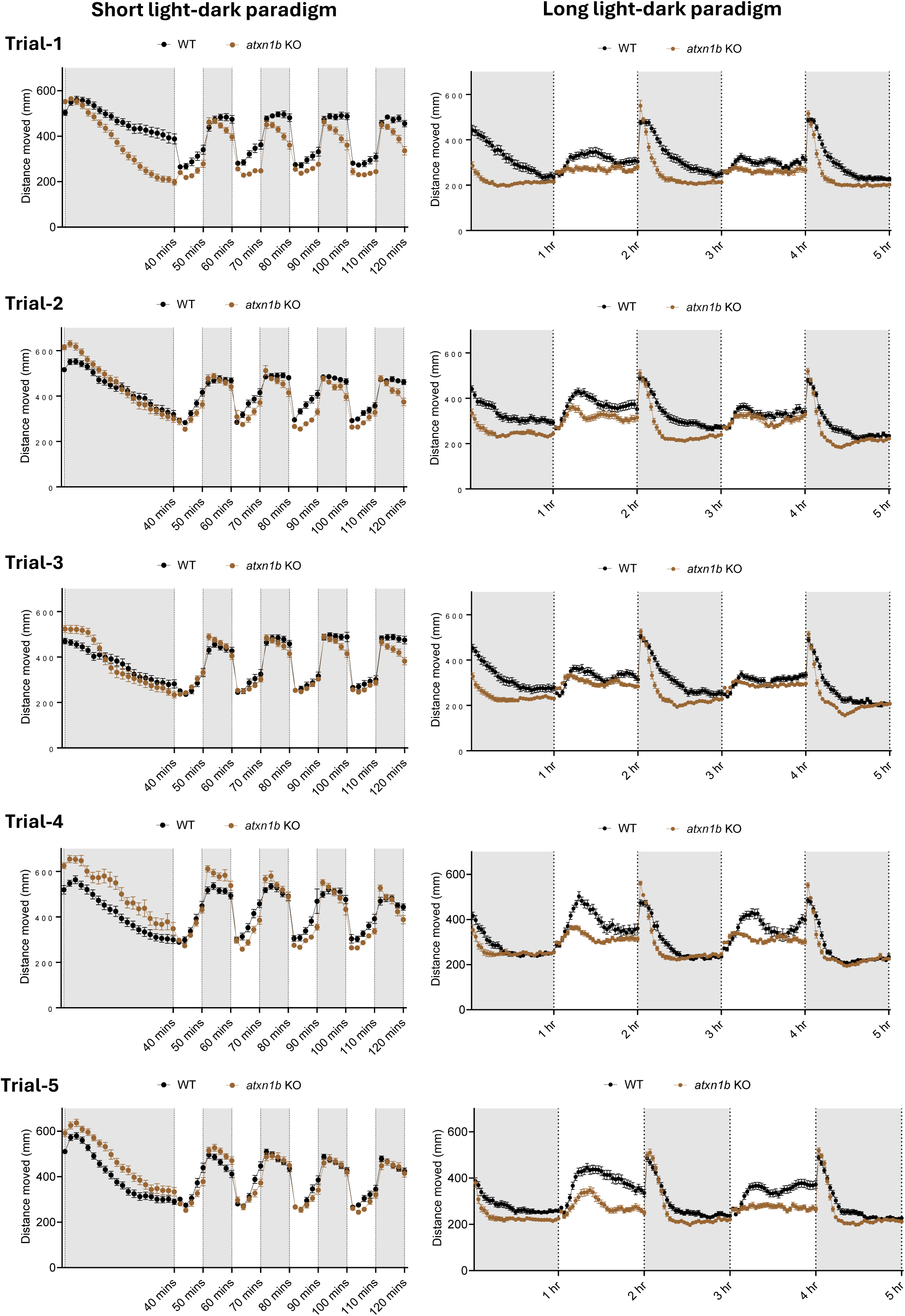
Assessment of locomotor activity of homozygous *atxn1b* KO larvae at 5 dpf. All five trials are shown. Left panel represents the short light-dark paradigm and the right panel represent long light-dark paradigm. Each trial was performed with 48 WT and 48 KO larvae.

**Supplementary Fig. 3.**
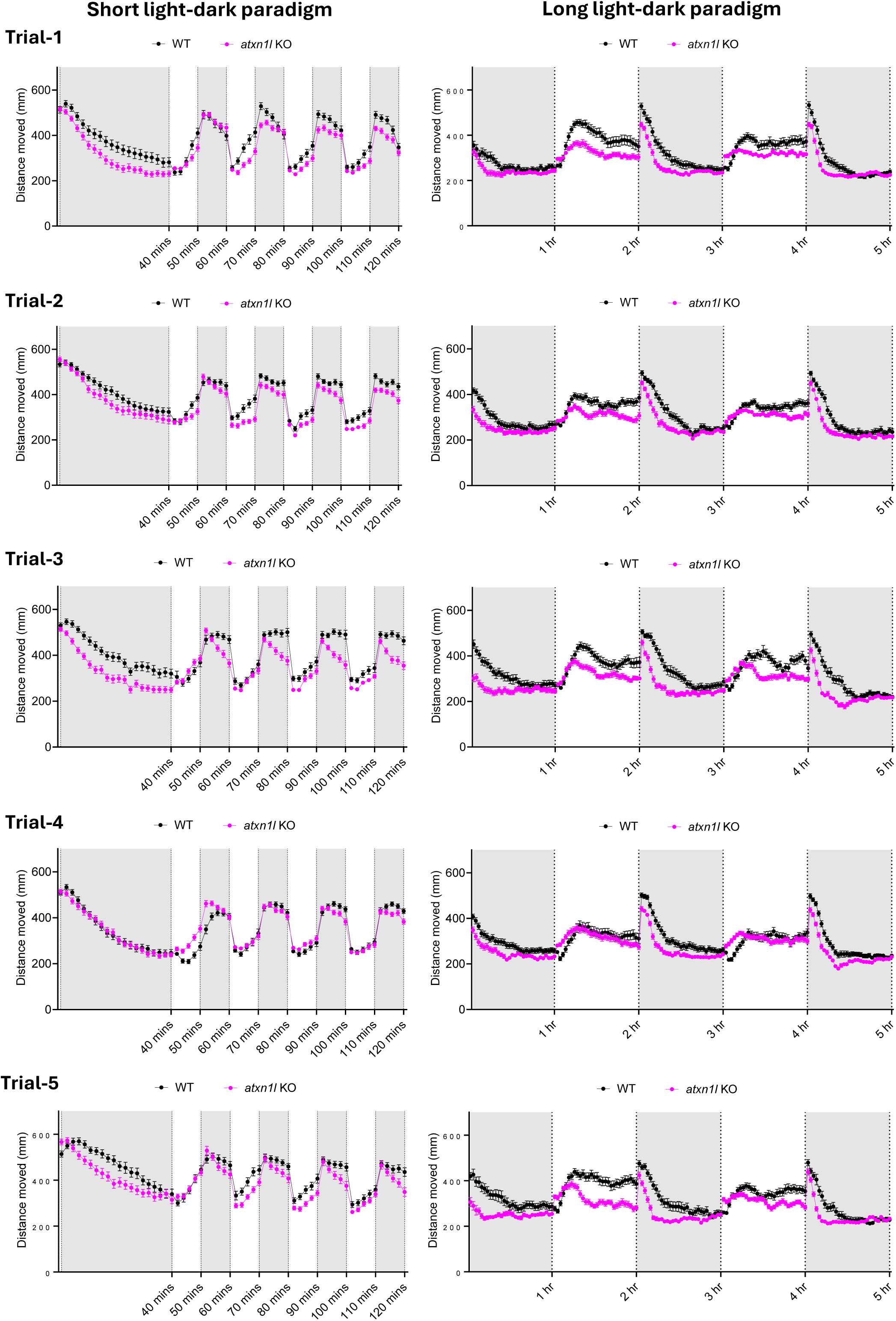
Assessment of locomotor activity of homozygous *atxn1l* KO larvae at 5 dpf. All five trials are shown. Left panel represents the short light-dark paradigm and the right panel represent long light-dark paradigm. Each trial was performed with 48 WT and 48 KO larvae.

**Supplementary Fig. 4:**
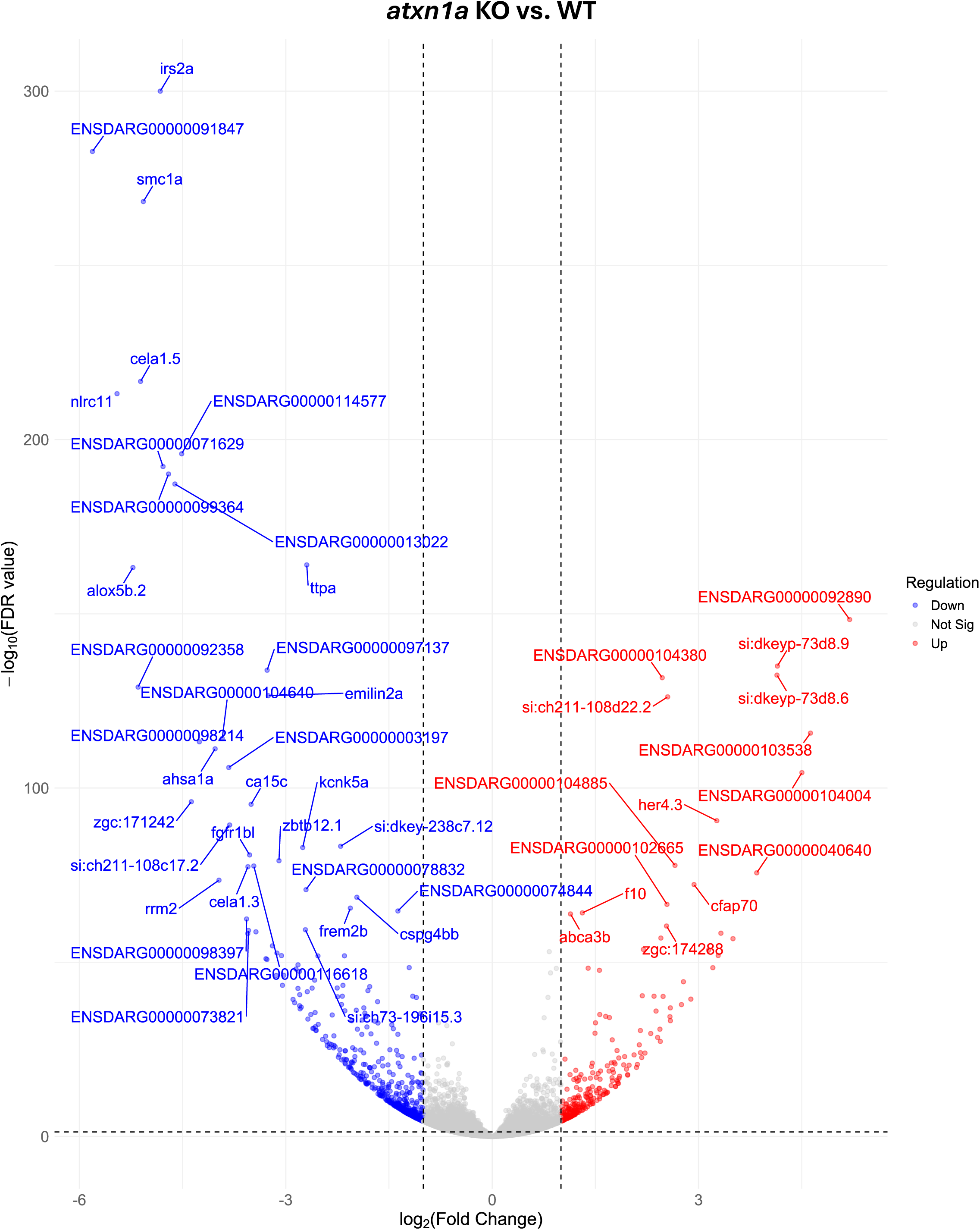
Volcano plot showing the result of differential expression analysis of *atxn1a* KO vs. WT larvae at 5 dpf. Top 50 significantly dysregulated genes are labeled.

**Supplementary Fig. 5:**
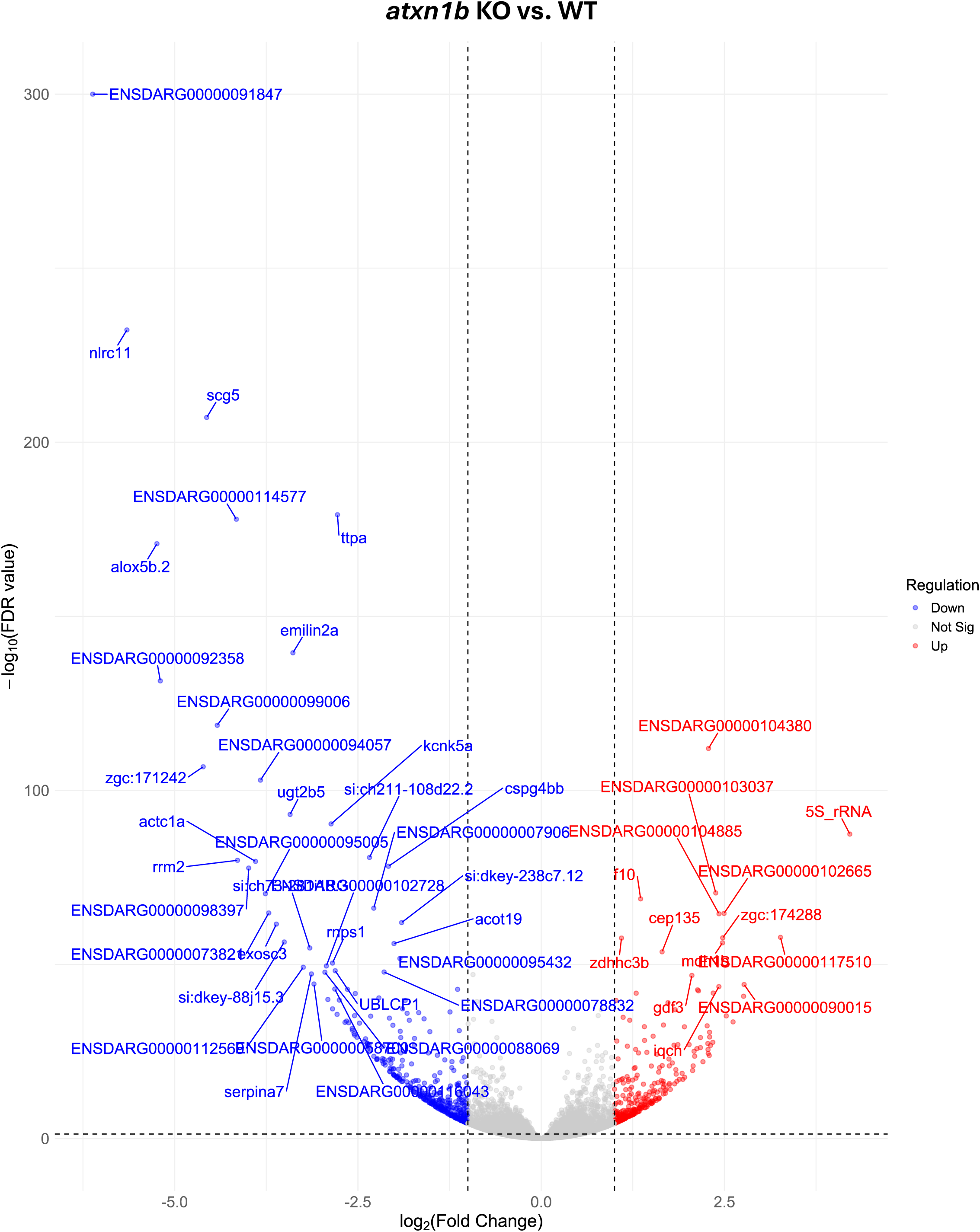
Volcano plot showing the result of differential expression analysis of *atxn1b* KO vs. WT larvae at 5 dpf. Top 50 significantly dysregulated genes are labeled.

**Supplementary Fig. 6:**
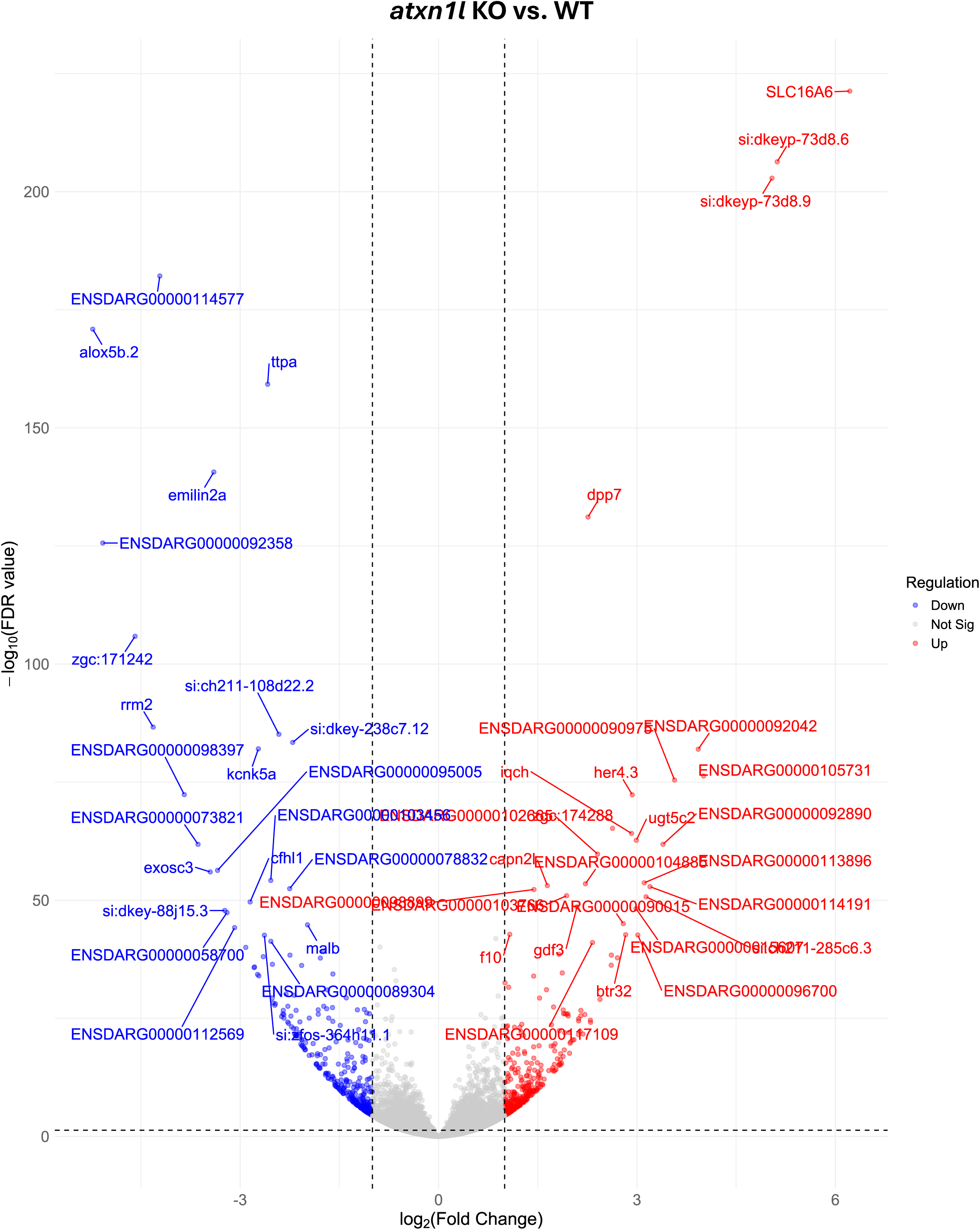
Volcano plot showing the result of differential expression analysis of *atxn1l* KO vs. WT larvae at 5 dpf. Top 50 significantly dysregulated genes are labeled.

**Supplementary Fig. 7:**
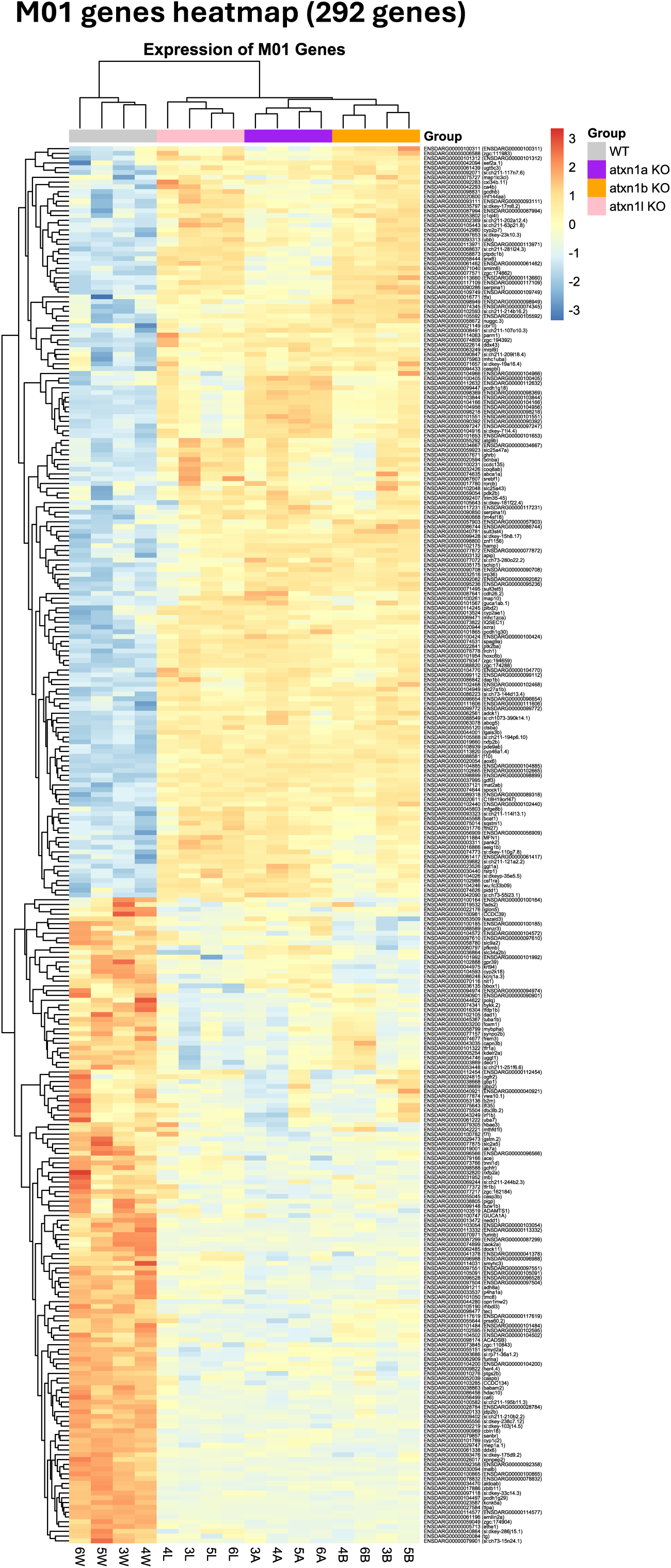
Heatmap showing the expression of M01 genes.

**Supplementary Fig. 8:**
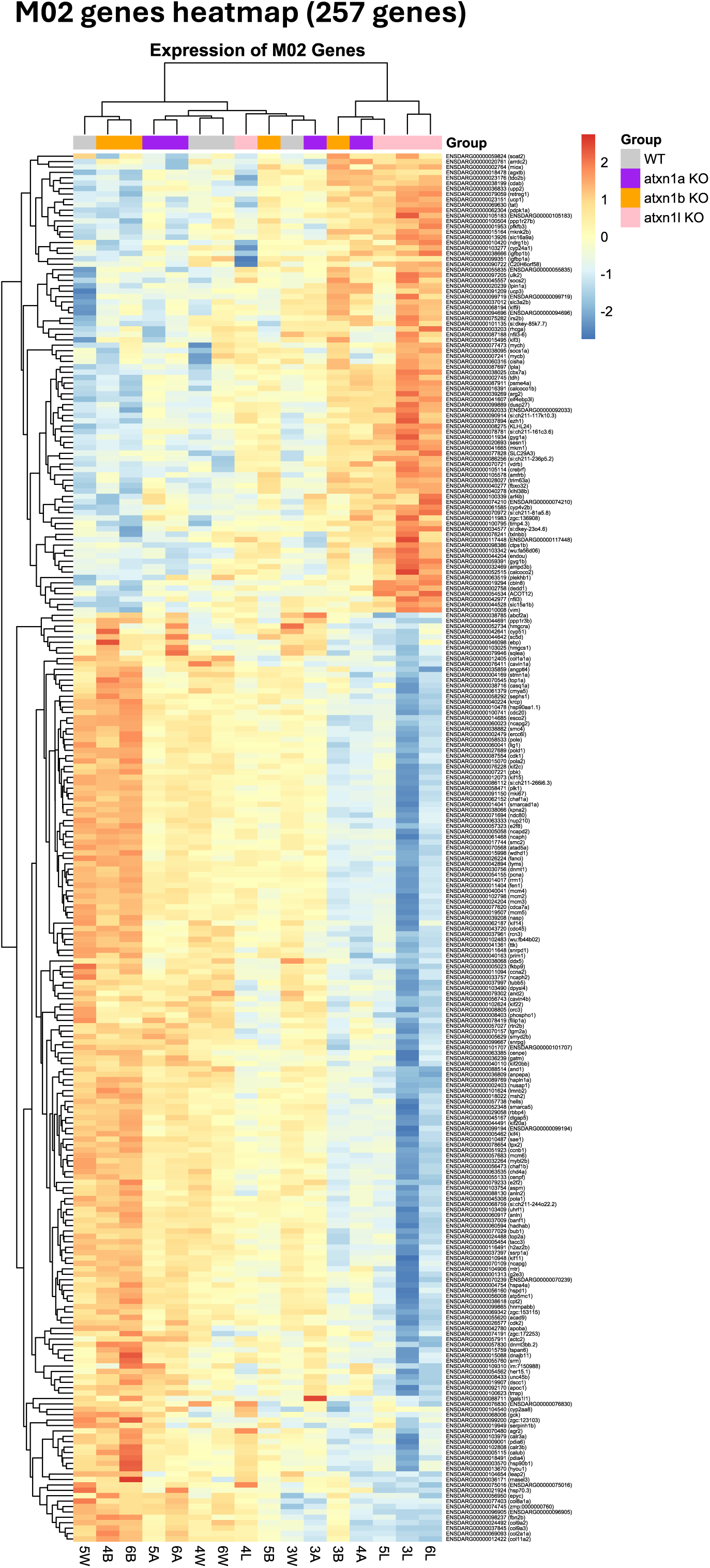
Heatmap showing the expression of M02 genes.

**Supplementary Fig. 9:**
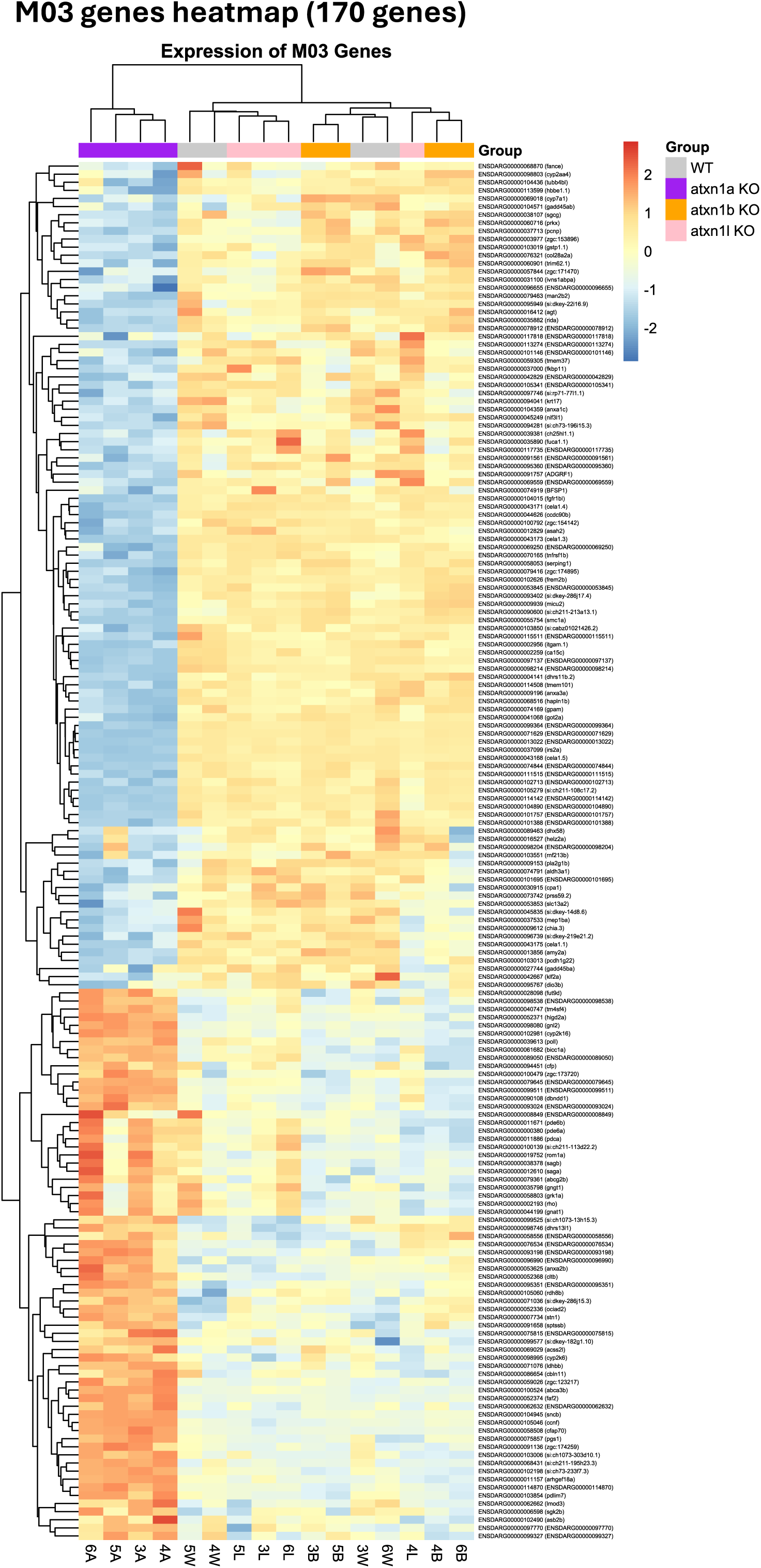
Heatmap showing the expression of M03 genes.

**Supplementary Fig. 10:**
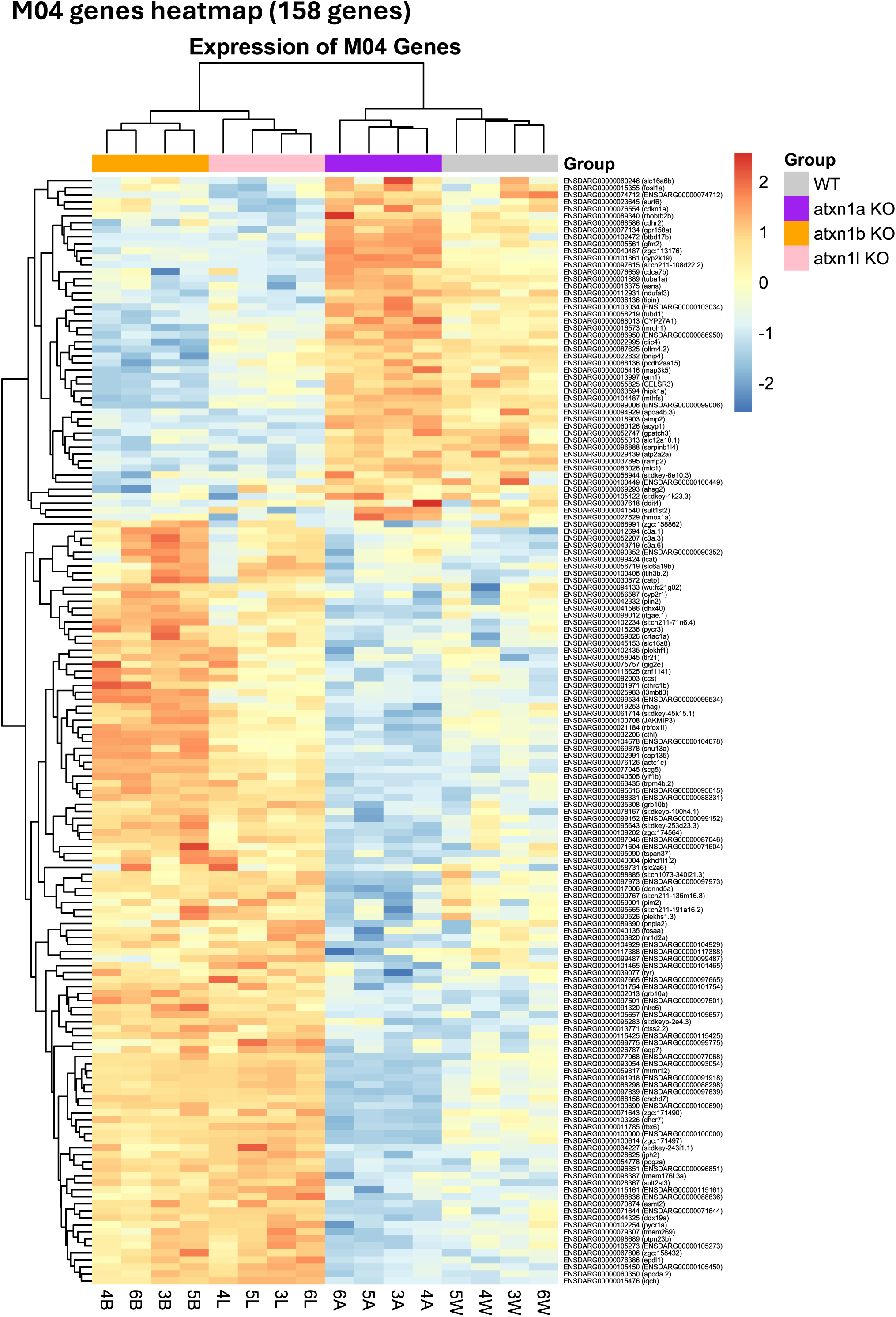
Heatmap showing the expression of M04 genes.

**Supplementary Fig. 11:**
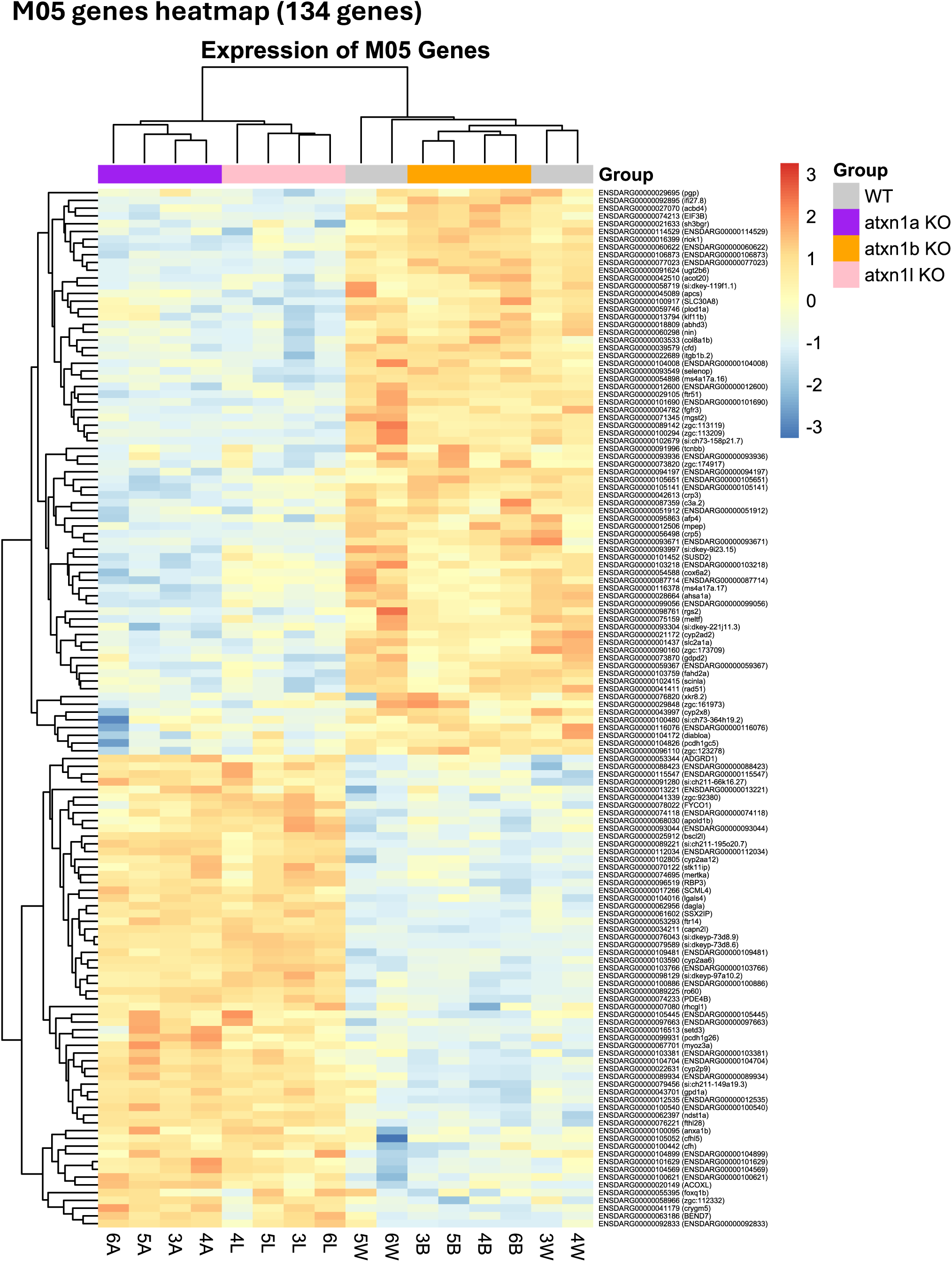
Heatmap showing the expression of M05 genes.

**Supplementary Fig. 12:**
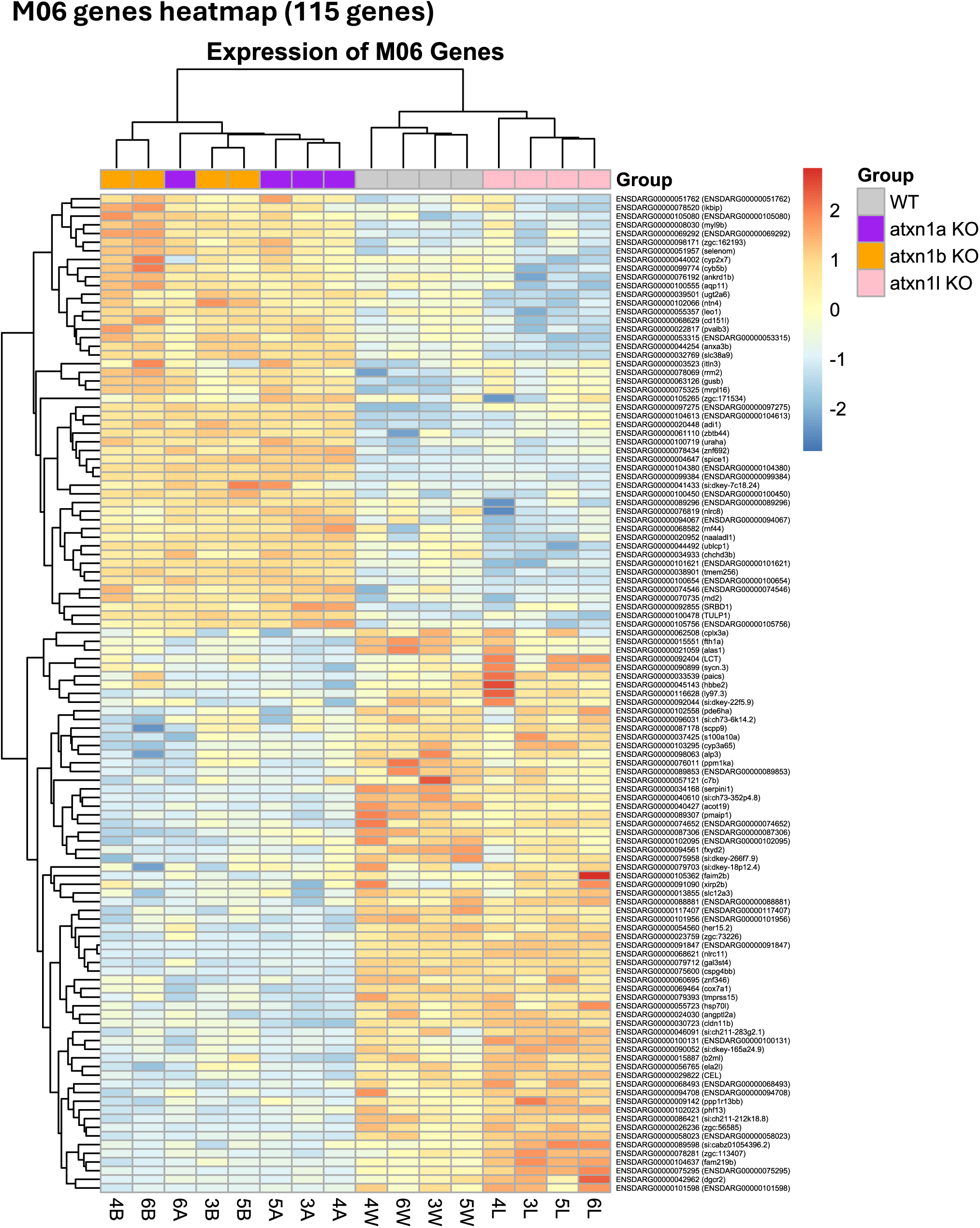
Heatmap showing the expression of M06 genes.

**Supplementary Fig. 13:**
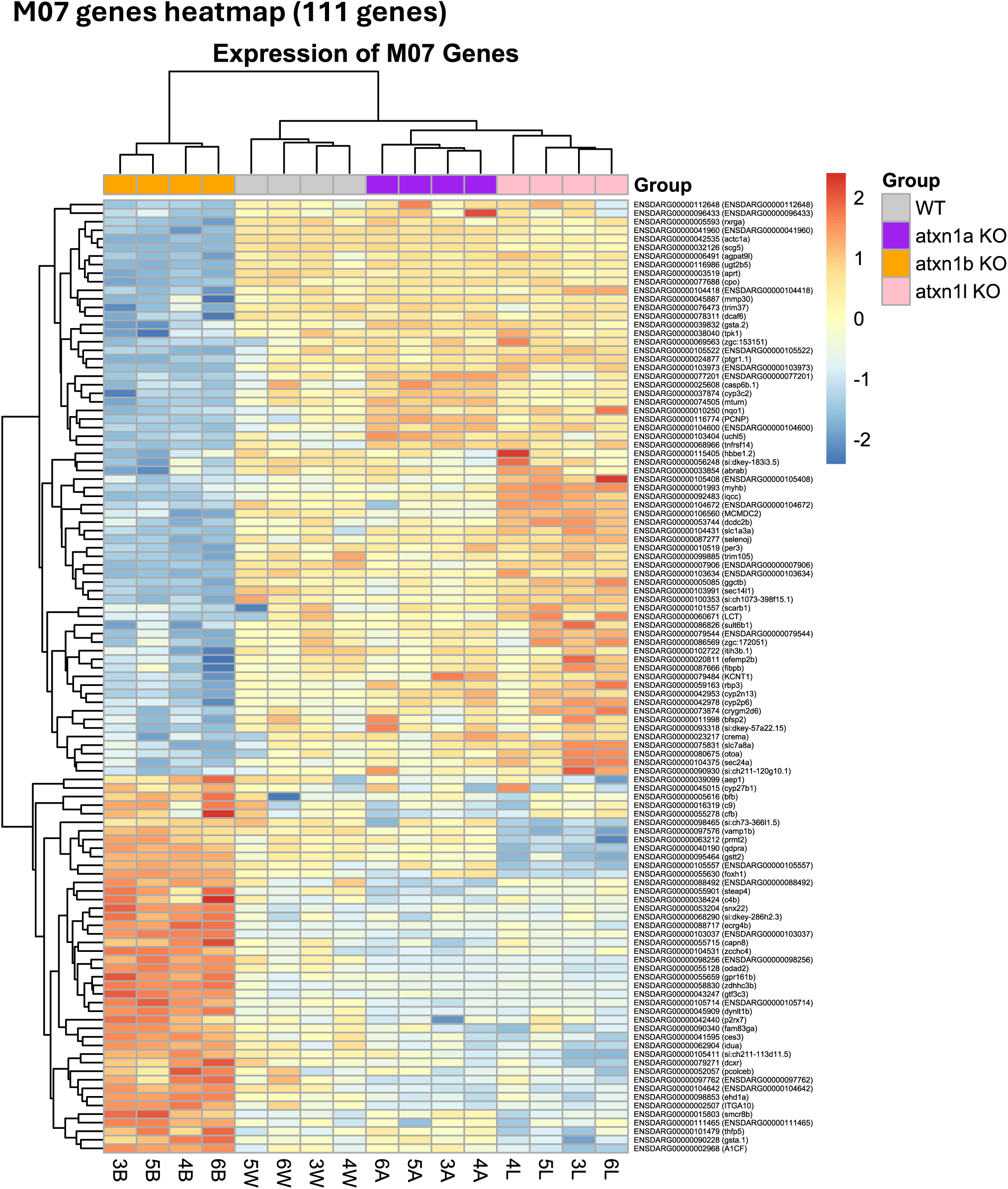
Heatmap showing the expression of M07 genes.

**Supplementary Fig. 14:**
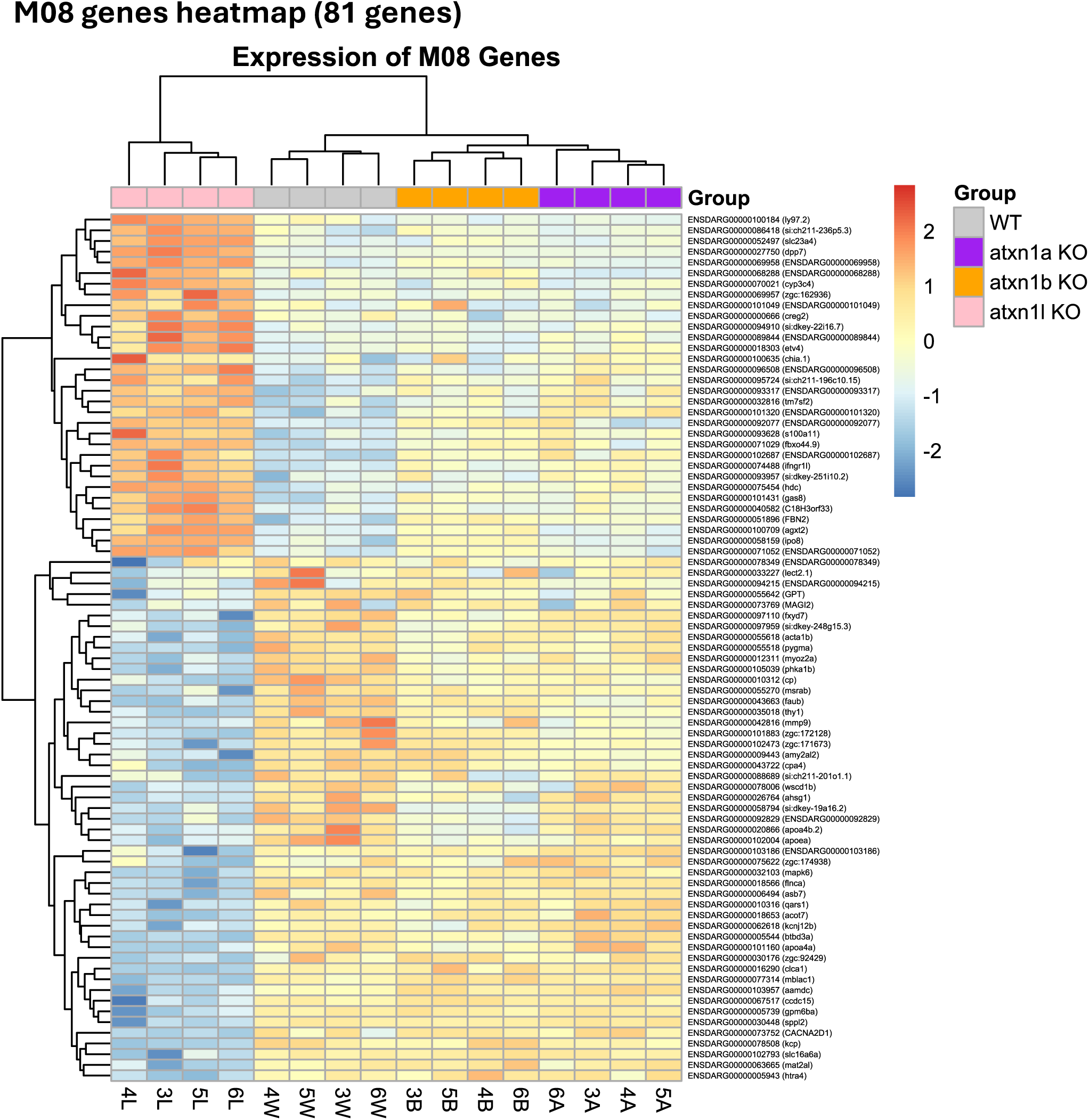
Heatmap showing the expression of M08 genes.

**Supplementary Fig. 15:**
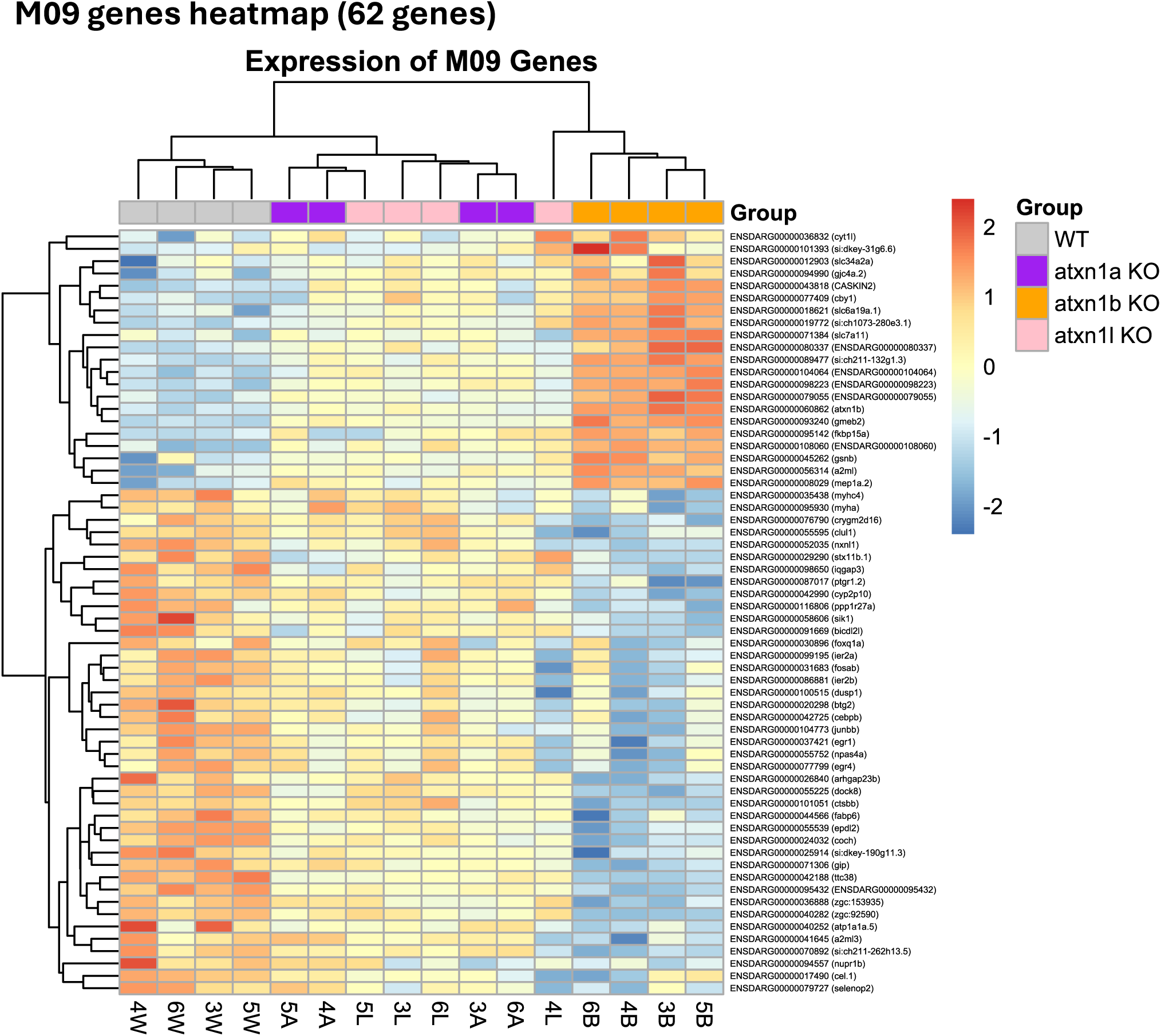
Heatmap showing the expression of M09 genes.

**Supplementary Fig. 16:**
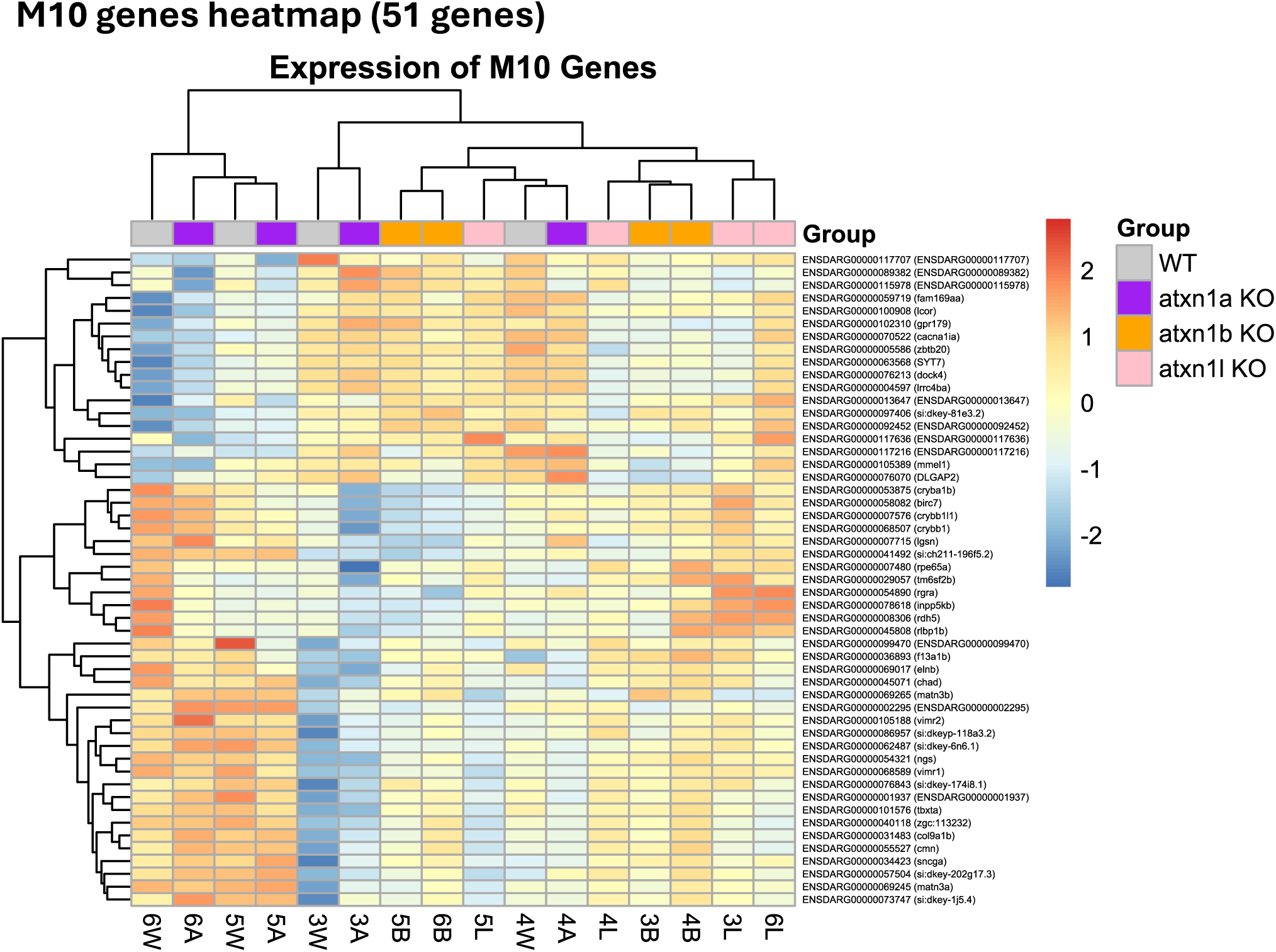
Heatmap showing the expression of M10 genes.

**Supplementary Fig. 17:**
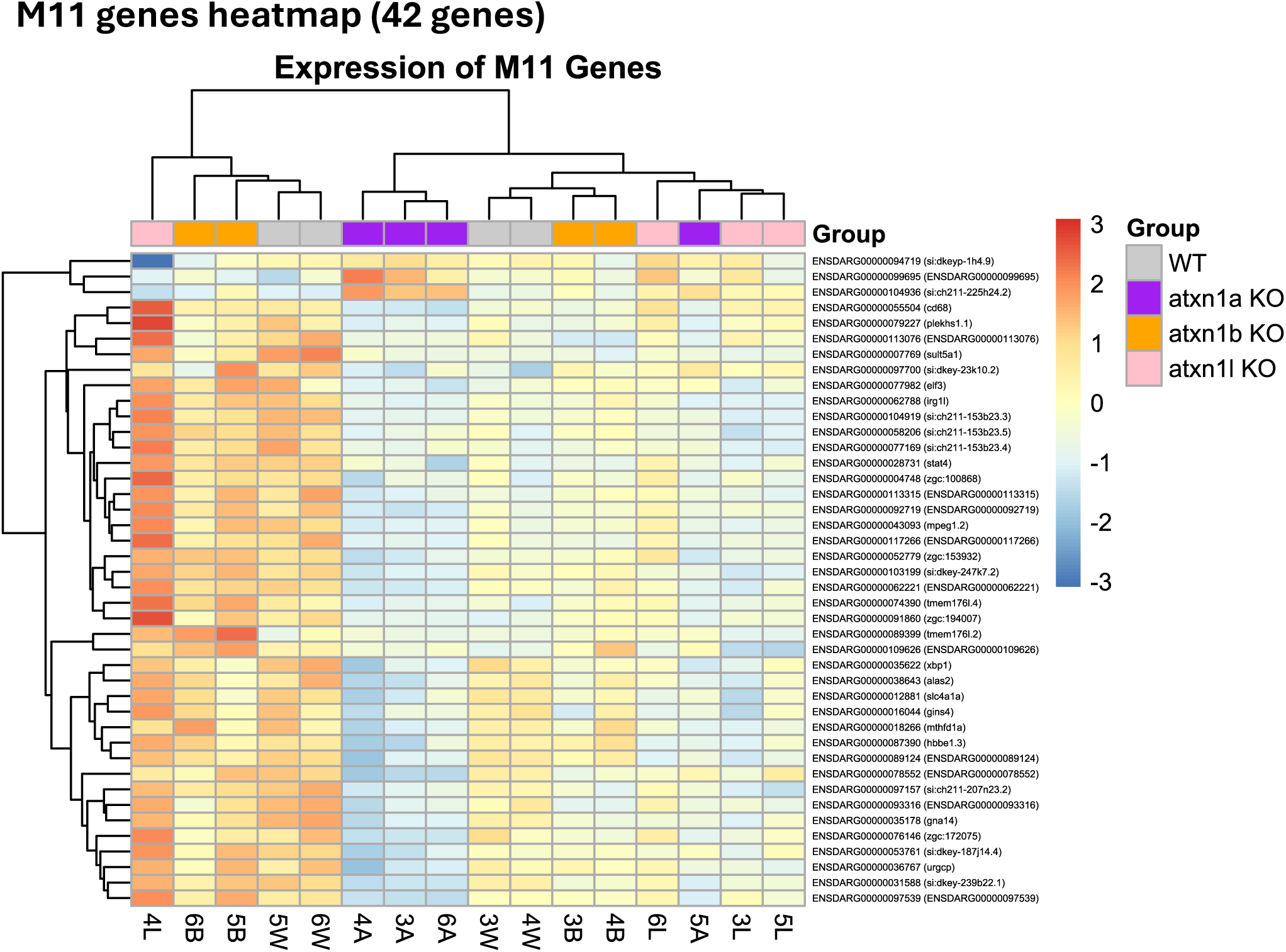
Heatmap showing the expression of M11 genes.

**Supplementary Fig. 18:**
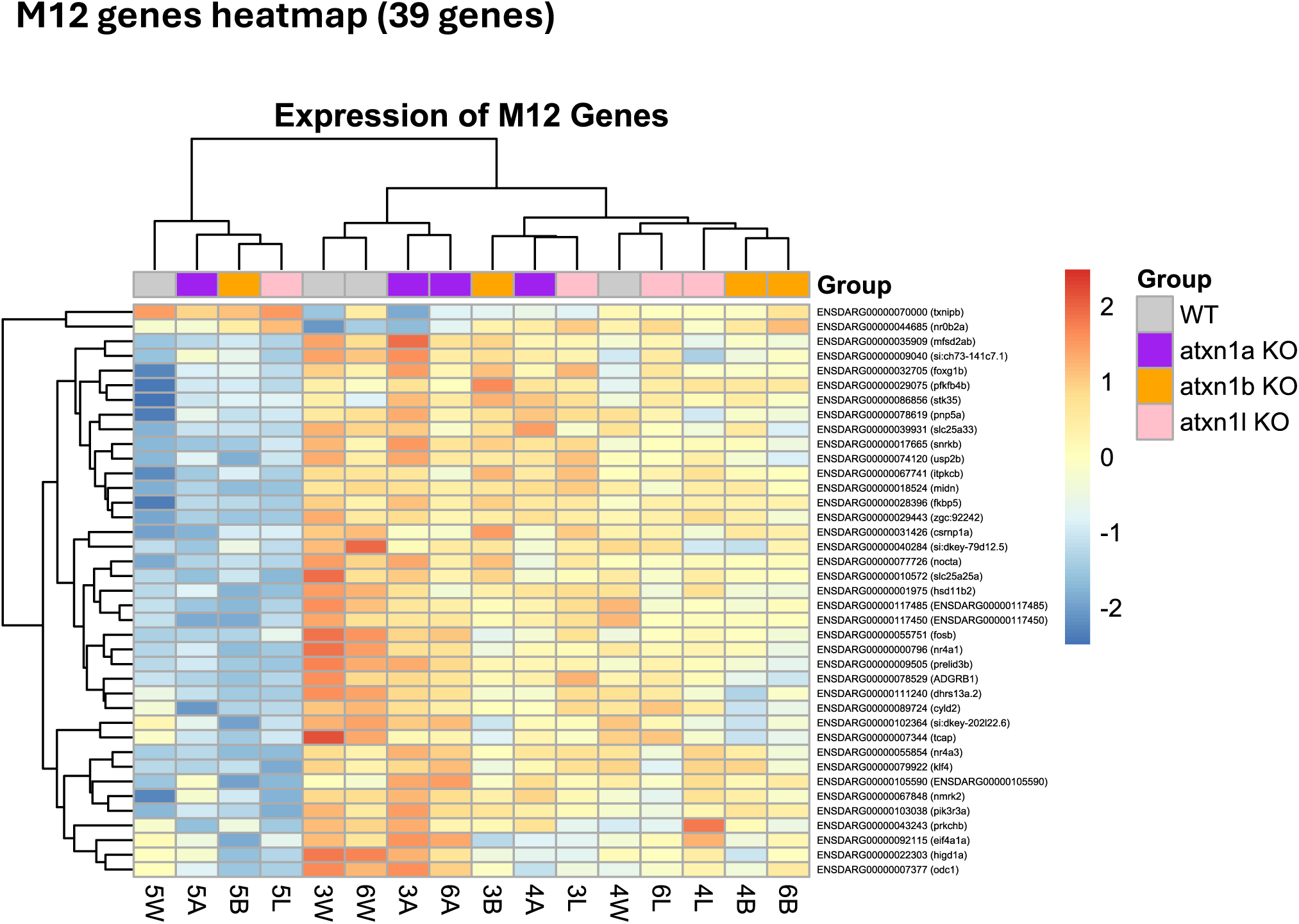
Heatmap showing the expression of M12 genes.

## References

Bansod, S., Kageyama, R., & Ohtsuka, T. (2017). Hes5 regulates the transition timing of neurogenesis and gliogenesis in mammalian neocortical development. Development, 144(17), 3156–3167. 10.1242/dev.147256

Bogdan, C. (2015). Nitric oxide synthase in innate and adaptive immunity: An update. Trends in Immunology, 36(3), 161–178. 10.1016/j.it.2015.01.003

Braccioli, L., Vervoort, S. J., Puma, G., Nijboer, C. H., & Coffer, P. J. (2018). SOX4 inhibits oligodendrocyte differentiation of embryonic neural stem cells in vitro by inducing Hes5 expression. Stem Cell Research, 33, 110–119. 10.1016/j.scr.2018.10.005

Carlson, K. M., Melcher, L., Lai, S., Zoghbi, H. Y., Clark, H. B., & Orr, H. T. (2009). Characterization of the Zebrafish *atxn1/axh* Gene Family. Journal of Neurogenetics, 23(3), 313–323. 10.1080/01677060802399976

Carrell, E. M., Keiser, M. S., Robbins, A. B., & Davidson, B. L. (2022). Combined overexpression of ATXN1L and mutant ATXN1 knockdown by AAV rescue motor phenotypes and gene signatures in SCA1 mice. Molecular Therapy - Methods & Clinical Development, 25, 333–343. 10.1016/j.omtm.2022.04.004

Carver, J. J., Denbrock, R. R., Martines, C. C., & Didonna, A. (2025). Conditional deletion of the multiple sclerosis susceptibility gene *ATXN1* identifies cell-autonomous effects in the B-cell compartment. The FEBS Journal, febs.70328. 10.1111/febs.70328

Chen, H., Dzitoyeva, S., & Manev, H. (2010). 5-Lipoxygenase in mouse cerebellar Purkinje cells. Neuroscience, 171(2), 383–389. 10.1016/j.neuroscience.2010.09.019

Cheng, C. W., Beech, D. J., & Wheatcroft, S. B. (2020). Advantages of CEMiTool for gene co-expression analysis of RNA-seq data. Computers in Biology and Medicine, 125, 103975. 10.1016/j.compbiomed.2020.103975

Chia, K., Klingseisen, A., Sieger, D., & Priller, J. (2022). Zebrafish as a model organism for neurodegenerative disease. Frontiers in Molecular Neuroscience, 15, 940484. 10.3389/fnmol.2022.940484

Colton, C. A., Vitek, M. P., Wink, D. A., Xu, Q., Cantillana, V., Previti, M. L., Van Nostrand, W. E., Weinberg, J. B., & Dawson, H. (2006). NO synthase 2 (*NOS2*) deletion promotes multiple pathologies in a mouse model of Alzheimer’s disease. Proceedings of the National Academy of Sciences, 103(34), 12867–12872. 10.1073/pnas.0601075103

Colton, C. A., Wilcock, D. M., Wink, D. A., Davis, J., Van Nostrand, W. E., & Vitek, M. P. (2008). The Effects of NOS2 Gene Deletion on Mice Expressing Mutated Human AβPP. Journal of Alzheimer’s Disease, 15(4), 571–587. 10.3233/JAD-2008-15405

Cote, R. H. (2021). Photoreceptor phosphodiesterase (PDE6): Activation and inactivation mechanisms during visual transduction in rods and cones. Pflügers Archiv - European Journal of Physiology, 473(9), 1377–1391. 10.1007/s00424-021-02562-x

Crespo-Barreto, J., Fryer, J. D., Shaw, C. A., Orr, H. T., & Zoghbi, H. Y. (2010). Partial Loss of Ataxin-1 Function Contributes to Transcriptional Dysregulation in Spinocerebellar Ataxia Type 1 Pathogenesis. PLoS Genetics, 6(7), e1001021. 10.1371/journal.pgen.1001021

De Sena Brandine, G., & Smith, A. D. (2021). Falco: High-speed FastQC emulation for quality control of sequencing data. F1000Research, 8, 1874. 10.12688/f1000research.21142.2

Didonna, A., Canto Puig, E., Ma, Q., Matsunaga, A., Ho, B., Caillier, S. J., Shams, H., Lee, N., Hauser, S. L., Tan, Q., Zamvil, S. S., & Oksenberg, J. R. (2020). Ataxin-1 regulates B cell function and the severity of autoimmune experimental encephalomyelitis. Proceedings of the National Academy of Sciences, 117(38), 23742–23750. 10.1073/pnas.2003798117

Dobin, A., Davis, C. A., Schlesinger, F., Drenkow, J., Zaleski, C., Jha, S., Batut, P., Chaisson, M., & Gingeras, T. R. (2013). STAR: Ultrafast universal RNA-seq aligner. Bioinformatics, 29(1), 15–21. 10.1093/bioinformatics/bts635

Elsaey, M. A., Namikawa, K., & Köster, R. W. (2021). Genetic Modeling of the Neurodegenerative Disease Spinocerebellar Ataxia Type 1 in Zebrafish. International Journal of Molecular Sciences, 22(14), 7351. 10.3390/ijms22147351

Gence, L., Fernezelian, D., Meilhac, O., Rastegar, S., Bascands, J., & Diotel, N. (2023). Insulin signaling promotes neurogenesis in the brain of adult zebrafish. Journal of Comparative Neurology, 531(17), 1812–1827. 10.1002/cne.25542

Golushko, N. I., Matrynov, D., Galstyan, D. S., Apukhtin, K. V., De Abreu, M. S., Yang, L., Stewart, A. M., & Kalueff, A. V. (2025). Understanding (and appreciating) behavioral complexity of zebrafish novel tank assays. Behavioural Processes, 230, 105230. 10.1016/j.beproc.2025.105230

Goold, R., Hubank, M., Hunt, A., Holton, J., Menon, R. P., Revesz, T., Pandolfo, M., & Matilla-Duenas, A. (2007). Down-regulation of the dopamine receptor D2 in mice lacking ataxin 1. Human Molecular Genetics, 16(17), 2122–2134. 10.1093/hmg/ddm162

Hoser, M., Baader, S. L., Bösl, M. R., Ihmer, A., Wegner, M., & Sock, E. (2007). Prolonged Glial Expression of Sox4 in the CNS Leads to Architectural Cerebellar Defects and Ataxia. The Journal of Neuroscience, 27(20), 5495–5505. 10.1523/JNEUROSCI.1384-07.2007

Huang, D. W., Sherman, B. T., & Lempicki, R. A. (2009). Systematic and integrative analysis of large gene lists using DAVID bioinformatics resources. Nature Protocols, 4(1), 44–57. 10.1038/nprot.2008.211

International Multiple Sclerosis Genetics Consortium, Patsopoulos, N. A., Baranzini, S. E., Santaniello, A., Shoostari, P., Cotsapas, C., Wong, G., Beecham, A. H., James, T., Replogle, J., Vlachos, I. S., McCabe, C., Pers, T. H., Brandes, A., White, C., Keenan, B., Cimpean, M., Winn, P., Panteliadis, I.-P., … De Jager, P. L. (2019). Multiple sclerosis genomic map implicates peripheral immune cells and microglia in susceptibility. Science, 365(6460), eaav7188. 10.1126/science.aav7188

Iova, O.-M., Marin, G.-E., Lazar, I., Stanescu, I., Dogaru, G., Nicula, C. A., & Bulboacă, A. E. (2023). Nitric Oxide/Nitric Oxide Synthase System in the Pathogenesis of Neurodegenerative Disorders—An Overview. Antioxidants, 12(3), 753. 10.3390/antiox12030753

Iribarne, M., & Masai, I. (2017). Neurotoxicity of cGMP in the vertebrate retina: From the initial research on *rd* mutant mice to zebrafish genetic approaches. Journal of Neurogenetics, 31(3), 88–101. 10.1080/01677063.2017.1358268

Keerthisinghe, P., Karim, A., Basak, A., & Orengo, J. P. (2026). Protocol for quantitative analysis of adult zebrafish swimming behavior using DeepLabCut. STAR Protocols, 7(1), 104374. 10.1016/j.xpro.2026.104374

Kerkhof, L. M. C., Warrenburg, B. P. C. V. D., Roon-Mom, W. M. C. V., & Buijsen, R. A. M. (2023). Therapeutic Strategies for Spinocerebellar Ataxia Type 1. Biomolecules, 13(5), 788. 10.3390/biom13050788

Kim, J., Han, J.-Y., Lee, Y., Kim, K., Choi, Y. P., Chae, S., & Hoe, H.-S. (2023). Genetic deletion of nitric oxide synthase 2 ameliorates Parkinson’s disease pathology and neuroinflammation in a transgenic mouse model of synucleinopathy. Molecular Brain, 16(1), 7. 10.1186/s13041-023-00996-1

Lam, Y. C., Bowman, A. B., Jafar-Nejad, P., Lim, J., Richman, R., Fryer, J. D., Hyun, E. D., Duvick, L. A., Orr, H. T., Botas, J., & Zoghbi, H. Y. (2006). ATAXIN-1 Interacts with the Repressor Capicua in Its Native Complex to Cause SCA1 Neuropathology. Cell, 127(7), 1335–1347. 10.1016/j.cell.2006.11.038

Lee, Y., Fryer, J. D., Kang, H., Crespo-Barreto, J., Bowman, A. B., Gao, Y., Kahle, J. J., Hong, J. S., Kheradmand, F., Orr, H. T., Finegold, M. J., & Zoghbi, H. Y. (2011). ATXN1 Protein Family and CIC Regulate Extracellular Matrix Remodeling and Lung Alveolarization. Developmental Cell, 21(4), 746–757. 10.1016/j.devcel.2011.08.017

Liao, Y., Smyth, G. K., & Shi, W. (2014). featureCounts: An efficient general purpose program for assigning sequence reads to genomic features. Bioinformatics, 30(7), 923–930. 10.1093/bioinformatics/btt656

Love, M. I., Huber, W., & Anders, S. (2014). Moderated estimation of fold change and dispersion for RNA-seq data with DESeq2. Genome Biology, 15(12), 550. 10.1186/s13059-014-0550-8

Ma, Q., & Didonna, A. (2023). Ataxin-1 controls the expression of specific noncoding RNAS in B cells upon autoimmune demyelination. Immunology & Cell Biology, 101(4), 358–367. 10.1111/imcb.12622

Martin, M. (2011). Cutadapt removes adapter sequences from high-throughput sequencing reads. EMBnet.Journal, 17(1), 10. 10.14806/ej.17.1.200

Mathis, A., Mamidanna, P., Cury, K. M., Abe, T., Murthy, V. N., Mathis, M. W., & Bethge, M. (2018). DeepLabCut: Markerless pose estimation of user-defined body parts with deep learning. Nature Neuroscience, 21(9), 1281–1289. 10.1038/s41593-018-0209-y

Matilla, A., Roberson, E. D., Banfi, S., Morales, J., Armstrong, D. L., Burright, E. N., Orr, H. T., Sweatt, J. D., Zoghbi, H. Y., & Matzuk, M. M. (1998). Mice Lacking Ataxin-1 Display Learning Deficits and Decreased Hippocampal Paired-Pulse Facilitation. The Journal of Neuroscience, 18(14), 5508–5516. 10.1523/JNEUROSCI.18-14-05508.1998

Mizutani, A., Wang, L., Rajan, H., Vig, P. J., Alaynick, W. A., Thaler, J. P., & Tsai, C. (2005). Boat, an AXH domain protein, suppresses the cytotoxicity of mutant ataxin-1. The EMBO Journal, 24(18), 3339–3351. 10.1038/sj.emboj.7600785

Montague, T. G., Cruz, J. M., Gagnon, J. A., Church, G. M., & Valen, E. (2014). CHOPCHOP: A CRISPR/Cas9 and TALEN web tool for genome editing. Nucleic Acids Research, 42(W1), W401–W407. 10.1093/nar/gku410

Muradov, H., Boyd, K. K., & Artemyev, N. O. (2010). Rod phosphodiesterase-6 PDE6A and PDE6B Subunits Are Enzymatically Equivalent. Journal of Biological Chemistry, 285(51), 39828–39834. 10.1074/jbc.M110.170068

Olmos, V., Gogia, N., Luttik, K., Haidery, F., & Lim, J. (2022). The extra-cerebellar effects of spinocerebellar ataxia type 1 (SCA1): Looking beyond the cerebellum. Cellular and Molecular Life Sciences, 79(8), 404. 10.1007/s00018-022-04419-7

Orr, H. T., Chung, M., Banfi, S., Kwiatkowski, T. J., Servadio, A., Beaudet, A. L., McCall, A. E., Duvick, L. A., Ranum, L. P. W., & Zoghbi, H. Y. (1993). Expansion of an unstable trinucleotide CAG repeat in spinocerebellar ataxia type 1. Nature Genetics, 4(3), 221–226. 10.1038/ng0793-221

Raudvere, U., Kolberg, L., Kuzmin, I., Arak, T., Adler, P., Peterson, H., & Vilo, J. (2019). g:Profiler: A web server for functional enrichment analysis and conversions of gene lists (2019 update). Nucleic Acids Research, 47(W1), W191–W198. 10.1093/nar/gkz369

Russo, P. S. T., Ferreira, G. R., Cardozo, L. E., Bürger, M. C., Arias-Carrasco, R., Maruyama, S. R., Hirata, T. D. C., Lima, D. S., Passos, F. M., Fukutani, K. F., Lever, M., Silva, J. S., Maracaja-Coutinho, V., & Nakaya, H. I. (2018). CEMiTool: A Bioconductor package for performing comprehensive modular co-expression analyses. BMC Bioinformatics, 19(1), 56. 10.1186/s12859-018-2053-1

Saleem, S., & Kannan, R. R. (2018). Zebrafish: An emerging real-time model system to study Alzheimer’s disease and neurospecific drug discovery. Cell Death Discovery, 4(1), 45. 10.1038/s41420-018-0109-7

Sánchez, I., Balagué, E., & Matilla-Dueñas, A. (2016). Ataxin-1 regulates the cerebellar bioenergetics proteome through the GSK3β-mTOR pathway which is altered in Spinocerebellar ataxia type 1 (SCA1). Human Molecular Genetics, 25(18), 4021–4040. 10.1093/hmg/ddw242

Sarasamma, S., Karim, A., & Orengo, J. P. (2023). Zebrafish Models of Rare Neurological Diseases like Spinocerebellar Ataxias (SCAs): Advantages and Limitations. Biology, 12(10), 1322. 10.3390/biology12101322

Schubert, M., Brazil, D. P., Burks, D. J., Kushner, J. A., Ye, J., Flint, C. L., Farhang-Fallah, J., Dikkes, P., Warot, X. M., Rio, C., Corfas, G., & White, M. F. (2003). Insulin Receptor Substrate-2 Deficiency Impairs Brain Growth and Promotes Tau Phosphorylation. The Journal of Neuroscience, 23(18), 7084–7092. 10.1523/JNEUROSCI.23-18-07084.2003

Schwartz, A. V., Sant, K. E., & George, U. Z. (2024). danRerLib: A Python package for zebrafish transcriptomics. Bioinformatics Advances, 4(1), vbae065. 10.1093/bioadv/vbae065

Sherman, B. T., Hao, M., Qiu, J., Jiao, X., Baseler, M. W., Lane, H. C., Imamichi, T., & Chang, W. (2022). DAVID: A web server for functional enrichment analysis and functional annotation of gene lists (2021 update). Nucleic Acids Research, 50(W1), W216–W221. 10.1093/nar/gkac194

Sonar, S. A., & Lal, G. (2019). The iNOS Activity During an Immune Response Controls the CNS Pathology in Experimental Autoimmune Encephalomyelitis. Frontiers in Immunology, 10, 710. 10.3389/fimmu.2019.00710

Srinivasan, S. R., & Shakkottai, V. G. (2019). Moving Towards Therapy in SCA1: Insights from Molecular Mechanisms, Identification of Novel Targets, and Planning for Human Trials. Neurotherapeutics, 16(4), 999–1008. 10.1007/s13311-019-00763-y

Srivastava, R., Eswar, K., Ramesh, S. S. R., Prajapati, A., Sonpipare, T., Basa, A., Gubige, M., Ponnapalli, S., Thatikonda, S., & Rengan, A. K. (2025). Zebrafish as a Versatile Model Organism: From Tanks to Treatment. MedComm – Future Medicine, 4(3), e70028. 10.1002/mef2.70028

Suh, J., Romano, D. M., Nitschke, L., Herrick, S. P., DiMarzio, B. A., Dzhala, V., Bae, J.-S., Oram, M. K., Zheng, Y., Hooli, B., Mullin, K., Gennarino, V. A., Wasco, W., Schmahmann, J. D., Albers, M. W., Zoghbi, H. Y., & Tanzi, R. E. (2019). Loss of Ataxin-1 Potentiates Alzheimer’s Pathogenesis by Elevating Cerebral BACE1 Transcription. Cell, 178(5), 1159–1175.e17. 10.1016/j.cell.2019.07.043

Sun, Q.-Y., Zhou, H.-H., & Mao, X.-Y. (2019). Emerging Roles of 5-Lipoxygenase Phosphorylation in Inflammation and Cell Death. Oxidative Medicine and Cellular Longevity, 2019, 1–9. 10.1155/2019/2749173

Szklarczyk, D., Kirsch, R., Koutrouli, M., Nastou, K., Mehryary, F., Hachilif, R., Gable, A. L., Fang, T., Doncheva, N. T., Pyysalo, S., Bork, P., Jensen, L. J., & von Mering, C. (2023). The STRING database in 2023: Protein–protein association networks and functional enrichment analyses for any sequenced genome of interest. Nucleic Acids Research, 51(D1), D638–D646. 10.1093/nar/gkac1000

Talukdar, G., Duvick, L., Yang, P., O’Callaghan, B., Fuchs, G. J., Cvetanovic, M., & Orr, H. T. (2025). An expanded polyglutamine in ATAXIN1 results in a loss-of-function that exacerbates severity of Multiple Sclerosis in an EAE mouse model. Journal of Neuroinflammation, 22(1), 127. 10.1186/s12974-025-03450-2

Tazelaar, G. H. P., Boeynaems, S., De Decker, M., Van Vugt, J. J. F. A., Kool, L., Goedee, H. S., McLaughlin, R. L., Sproviero, W., Iacoangeli, A., Moisse, M., Jacquemyn, M., Daelemans, D., Dekker, A. M., Van Der Spek, R. A., Westeneng, H.-J., Kenna, K. P., Assialioui, A., Da Silva, N., Project MinE ALS Sequencing Consortium, … Van Es, M. A. (2020). *ATXN1* repeat expansions confer risk for amyotrophic lateral sclerosis and contribute to TDP-43 mislocalization. Brain Communications, 2(2), fcaa064. 10.1093/braincomms/fcaa064

Tejwani, L., & Lim, J. (2020). Pathogenic mechanisms underlying spinocerebellar ataxia type 1. Cellular and Molecular Life Sciences, 77(20), 4015–4029. 10.1007/s00018-020-03520-z

The Galaxy Community, Abueg, L. A. L., Afgan, E., Allart, O., Awan, A. H., Bacon, W. A., Baker, D., Bassetti, M., Batut, B., Bernt, M., Blankenberg, D., Bombarely, A., Bretaudeau, A., Bromhead, C. J., Burke, M. L., Capon, P. K., Čech, M., Chavero-Díez, M., Chilton, J. M., … Zoabi, R. (2024). The Galaxy platform for accessible, reproducible, and collaborative data analyses: 2024 update. Nucleic Acids Research, 52(W1), W83–W94. 10.1093/nar/gkae410

Tuz-Sasik, M. U., Boije, H., & Manuel, R. (2022). Characterization of locomotor phenotypes in zebrafish larvae requires testing under both light and dark conditions. PLOS ONE, 17(4), e0266491. 10.1371/journal.pone.0266491

Vauti, F., Vögele, V., Deppe, I., Hahnenstein, S. T., & Köster, R. W. (2021). Structural Analysis and Spatiotemporal Expression of Atxn1 Genes in Zebrafish Embryos and Larvae. International Journal of Molecular Sciences, 22(21), 11348. 10.3390/ijms222111348

Wang, P., Luo, L., & Chen, J. (2024). Her4.3+ radial glial cells maintain the brain vascular network through activation of Wnt signaling. Journal of Biological Chemistry, 300(8), 107570. 10.1016/j.jbc.2024.107570

Westerfield, M. (2000). The Zebrafish Book. A Guide for the Laboratory Use of Zebrafish (Danio Rerio) (4th ed.). Univ. of Oregon Press, Eugene.

Wisenden, B. D., Paulson, D. C., & Orr, M. (2022). Zebrafish embryos hatch early in response to chemical and mechanical indicators of predation risk, resulting in underdeveloped swimming ability of hatchling larvae. Biology Open, 11(12), bio059229. 10.1242/bio.059229

Wu, L., Xue, R., Chen, J., & Xu, J. (2022). Dock8 deficiency attenuates microglia colonization in early zebrafish larvae. Cell Death Discovery, 8(1), 366. 10.1038/s41420-022-01155-6

Zoghbi, H. Y., & Orr, H. T. (1995). Spinocerebellar ataxia type 1. Seminars in Cell Biology, 6(1), 29–35. 10.1016/1043-4682(95)90012-8

